# Genome-wide annotation and analyses of bifunctional genes in the human genome

**DOI:** 10.64898/2026.01.28.702170

**Authors:** Janki Insan, Manoj B Menon, Sonam Dhamija

## Abstract

Conventional gene annotation pipelines classify eukaryotic genes into protein-coding and non-coding. Alternative splicing may generate non-coding transcript variants from protein-coding genes, that are expressed in tissue- or disease- specific manner. We and others have described the genes which transcribe both coding and non-coding transcripts as ‘bifunctional genes’. Here we present a genome-wide analyses of bifunctional genes and reannotate the genes in the human genome reference assembly into coding, non-coding and bifunctional. We identify over 4000 bifunctional genes in the human genome, constituting approximately 10% of the transcribed genes, and present evidence that these genes are conserved in evolution and their number correlate well with genome size and complexity. These genes are enriched in gene sets involved in vesicular transport, autophagy, RNA/DNA binding, glycosylation and splicing. By monitoring the expression of non-coding exons in long-read sequencing datasets and by quantitative RT-PCR, we provide evidence for the expression of non-coding variants from bifunctional genes. The ncRNA transcripts from these genes might have similar or different roles from their cognate mRNA counterparts. They may act as miRNA sponges or harbour non-canonical open-reading frames that encode microproteins, while also competing for binding with RNA-binding proteins. We present evidence for establishing potential biological functions of bifunctional genes and summarise the findings in a searchable database. Further studies and functional characterization focused on this special group of genes may reveal interesting gene regulatory mechanisms relevant to physiology and pathology.

## INTRODUCTION

Advances in next generation and long-read sequencing technologies are leading to the discovery of novel transcript variants of human genes at an unprecedented rate.^1–6^ Novel not-in-catalogue transcripts discovered from such studies are often products of alternative splicing or variant-associated mis-splicing, are hypothesized to contribute to gene regulation, protein diversity, and have been explored for their potential to generate cancer-specific neoantigens.^5,7,8^ Collation of these novel transcripts has led to an increase in the total number of annotated transcripts in the GENCODE project^9^, from 229,580 transcripts in release 35 (March 2020) to 507,365 transcripts in release 49 (Sept 2025). This has also led to a significant increase in the number of transcripts annotated as ‘non-coding’ or ‘processed transcripts’.^9–11^ These transcripts do not have any annotated open reading frames (ORFs) and are often thought to be targets of non-sense mediated decay.^12,13^ However, growing evidence shows that the expression of these annotated ‘non-coding’ transcripts is regulated in a context- or disease-specific manner.^14–18^ From UV-induced DNA repair to epithelial-to-mesenchymal transition, diverse roles of ‘non-coding’ isoforms have been coming to the fore.^14,17^ On the other hand, dozens of lncRNAs harbour non-canonical ORFs and are reported to undergo translation to give rise to microproteins (miPs) and peptideins.^19–24^ As a result of this, genes earlier described as purely non-coding are now being bio-typed as ‘protein-coding’.^25–28^ Moreover, the wide-spread discovery of circular RNAs from protein-coding genes further adds to mammalian transcriptome complexity.^29,30^ Thus, despite in-depth characterization of human transcriptomes facilitated by advances in sequencing technologies, current gene annotation strategies inadequately annotate genes only as either ‘protein-coding’ and ‘non-coding’ genes.

As our understanding of transcript diversity and functionality from individual gene loci evolve, re-examining current annotations to account for these new discoveries has become imperative. We and others have proposed the use of the term ‘bifunctional genes’ while referring to genes which transcribe both coding and non-coding transcripts.^31,32^ As proposed earlier, non-coding splice variants of protein-coding genes can have an array of roles. They can interfere with the protein function downstream through translation of small non-canonical ORFs.^33^ Similar regulation of protein function by shorter splice-isoforms have been reported for genes like ZBP1^34^, PRMT5^35^ and CFLAR^36^ among others. The non-coding splice variants can also act as competing endogenous RNA (ceRNA) for miRNAs or compete with protein-coding transcripts for binding RNA-binding proteins.^31^ For example, *lncRNA-PNUTS*, a splice-variant of the *PPP1R10* gene was shown to be induced by TGFβ and function as a ceRNA to sponge miR-205, facilitating the expression of EMT transcription factor ZEB1.^14^

In the present study, we performed a genome-wide analyses to identify bifunctional genes in the human genome. Our reannotation of human RefSeq genome assembly revealed that over 20% of all protein-coding genes in the human genome have at least one predicted or validated non-coding splice variant. We also investigated the chromosomal distribution, strand distribution as well as functional enrichment of human bifunctional genes. Furthermore, we explored whether bifunctionality is tied to organismal complexity and evolution. In addition to providing supporting data for expression of both coding and non-coding variants from bifunctional genes based on reanalyses of short and long-read sequencing datasets, we also experimentally verified the expression of non-coding transcript variants from select set of bifunctional genes across cells from different lineages. In an effort to investigate potential functions of non-coding splice variants from bifunctional genes, we explored their possible role as ceRNAs and smORF-encoding transcripts. To facilitate further research on bifunctional genes, our findings are made available in the form of a searchable database.

## RESULTS

### Over 10% of all transcribed genes in the human genome are bifunctional

Annotated human genome sequence with aligned non-coding and coding transcript information is available from both the GENCODE project^9^ and the NCBI Reference Sequence Database (RefSeq)^10^. While the GENCODE project compiles all possible transcripts including those without sufficient experimental proof to provide comprehensive coverage, the NCBI RefSeq assemblies provide better curated data minimizing redundancy and increasing stability. To identify bifunctional genes in the human genome, we analysed RNA transcripts from the fully annotated NCBI RefSeq assembly (GRCh38.p14, Genome Reference Consortium Human Build 38 patch release 14)^10^. The GRCh38.p14 RefSeq assembly lists 67,356 genes, of which 42,987 genes harbour one or more transcripts (Figure 1A and Table S1). Interestingly, a significant number of genes in GRCh38.p14 are ’pseudogenes’ with no transcripts annotated. Among the genes harbouring transcripts, 4,387 genes have at least one coding and one non-coding transcript, including both validated and predicted transcripts, which we refer to as ‘bifunctional’ genes. If our proposed annotation of genes is contrasted against the current biotypes provided by the NCBI, over 20% of genes with the ‘protein-coding’ biotype are bifunctional. In addition, we also reclassified 19 non-coding genes as bifunctional, based on the presence of annotated predicted coding transcripts (Figure 1B and Data S1). The schematic in Figure 1C is an example for a bifunctional gene locus and illustrates how a bifunctional gene can transcribe an array of coding and non-coding transcripts with complex regulatory potential. Even in the more conventional RefSeq annotations, there are a large number of predicted transcripts, and majority of them are non-coding variants. This raises the legitimate question whether these non-verified transcripts are leading to an overrepresentation of bifunctional genes in our analyses. To test this, we repeated bifunctional gene annotation, considering only validated coding and non-coding transcripts. Of the 29820 total genes with only validated transcripts, 3114 genes were annotated as bifunctional (Figure S1 and Data S2). Thus, independent of the presence or absence of predicted variants, approximately 10% of the RefSeq genes are bifunctional (Figure S1B).

**Figure 1.**
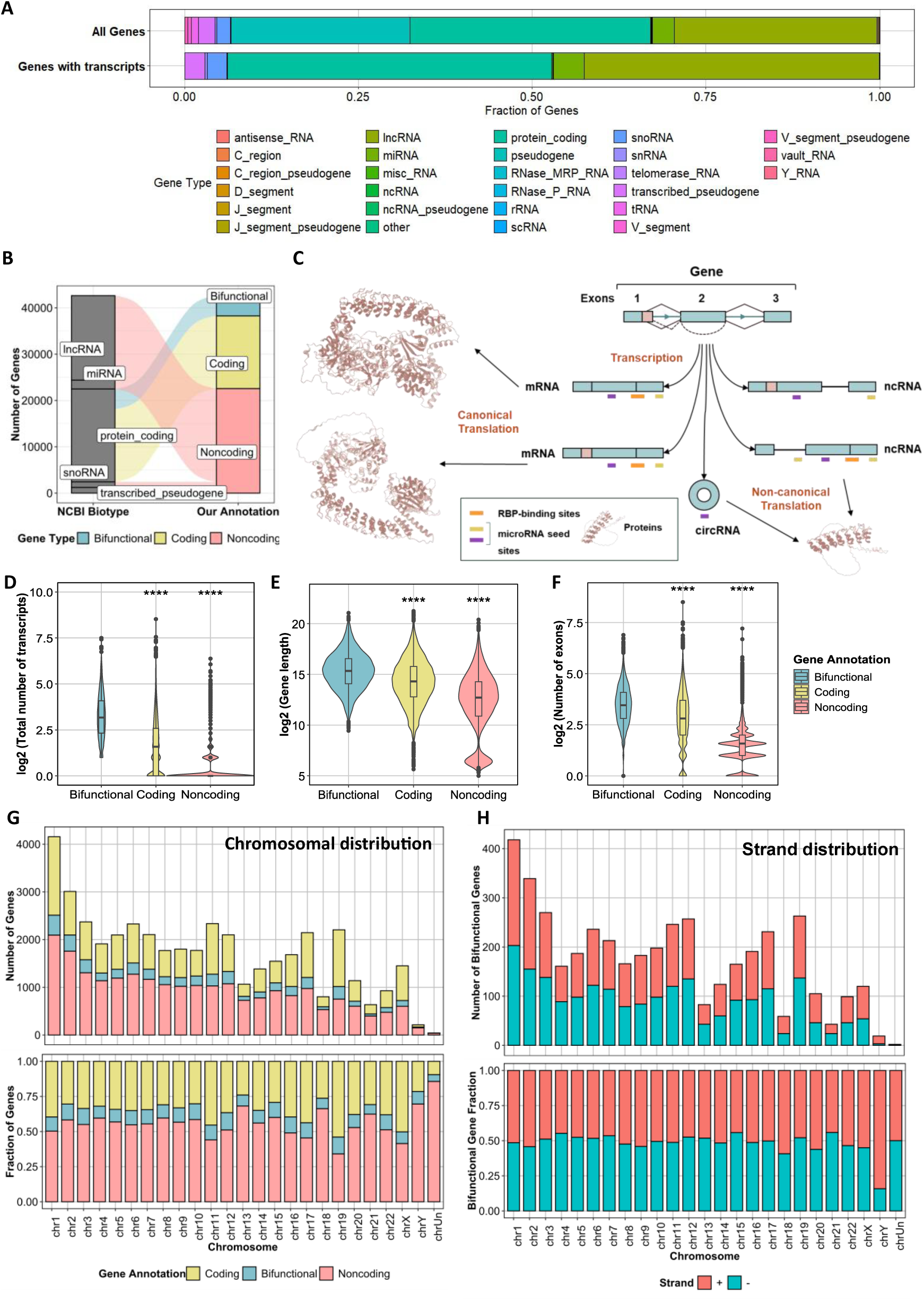
Identification and genome-wide analyses of bifunctional genes. **(A)** A distribution of all annotated genes versus transcribed genes from NCBI RefSeq, **(B)** Alluvial plot showing the distribution of NCBI biotypes versus our annotation based on the transcription products of each gene. **(C)** Depiction of a bifunctional gene locus showing how a single gene can generate both coding and non-coding RNA isoforms with distinct regulatory features and functions. **(D-F)** Differential analyses of number of transcripts (D), gene lengths (E) and number of exons in the longest transcript for bifunctional genes versus purely coding and non-coding genes (Pairwise comparisons were made using t-test, **** denotes p ≤ 0.0001, *** denotes p ≤ 0.001, ** denotes p ≤ 0.01 and * is used for p ≤ 0.05 respectively). **(G)** Chromosomal distribution of bifunctional genes versus genes with only mRNA and ncRNA transcripts. **(H)** Distribution of bifunctional genes on different strands.

To verify our findings, we extended our analyses to the independent RefSeq telomere-to-telomere genome assembly - T2T-CHM13v2.0 (Telomere-to-Telomere assembly of the CHM13 cell line, with chrY from NA24385). A similar analysis as in Figure 1B revealed 4,333 bifunctional genes in the T2T-CHM13v2.0 assembly. To further assess the consistency of our biotype annotations between the genome assemblies, we compared the biotypes of individual genes across the assemblies using their gene symbols (Figure S2A). Such an analysis revealed 4,186 genes which were consistently found to be bifunctional (Figure S2B, and Data S3). Similar results were obtained for other biotypes across both the assemblies (Figure S2C-D).

As most ‘bifunctional genes’ show complex patterns of alternative splicing, we then explored whether characteristics such as gene length, exon and transcript numbers correlate with bifunctionality. We found that bifunctional genes have longer gene lengths, higher exon numbers, and give rise to more transcripts than the pure coding and non-coding gene counterparts (Pairwise comparisons of bifunctional genes with their coding and non-coding counterparts gave a p<0.0001 for all comparisons using t-test and moderate to large values for Cohen’s d effect sizes; detailed values in Table S2) (Figure 1D-F). Since bifunctionality arises due to higher rates of alternative splicing events, our findings are in line with previous report that longer genes generate more transcript variants.^38^ We then looked at the chromosomal distribution of bifunctional genes across the genome. There was no significant enrichment of bifunctional genes in any of the chromosomes and the distribution was similar to that of coding and non-coding genes (Figure 1G and Table S3). When looking at the strand-specific distribution, we found that bifunctional genes are present on both strands, except for a minor enrichment on the positive strand in case of chromosomes 18 and Y (Figure 1H and Table S3). It is unclear whether this is due to the fact that these chromosomes display the lowest gene densities amongst human chromosomes. Similar results were obtained when this analysis was restricted to the annotations consisting of only the validated RefSeq transcripts (Figure S1C-F).

### The number of bifunctional genes correlates with genome size and complexity

Next, we examined whether organismal complexity is linked to bifunctionality of genes. To understand how bifunctional genes emerged in evolution, we analysed RefSeq assemblies of key model organisms with varying levels of organismal complexity (Data S3). We found that the number of bifunctional genes generally does increase with the increasing genome size, and an increase in the total number of transcribed genes, indicating that bifunctionality might be contributing to or be an outcome of increasing genome complexity (Figure 2A and 2B).^39,40^ *Caenorhabditis elegans* (worm) seems to be an exception to this observation, with large number of genes of all biotypes. On the other hand, the *Pan troglodytes* (chimpanzee) genome has fewer bifunctional genes than those found in *Mus musculus* (mouse). This could be attributed to the fact that the chimpanzee genome was most recently sequenced^41^ and have not been subjected to extensive transcript annotations compared to mouse or human genomes. Moreover, in the evolutionary timeline, the number of bifunctional genes in each organism more closely mirrors the increase in the genome size than the total number of genes (Figure 2B). A linear regression analysis of the same also reveals that the number of bifunctional genes in an organism strongly correlates with their overall genome size (R=0.78, p=0.008; Figure S3A-B).

**Figure 2.**
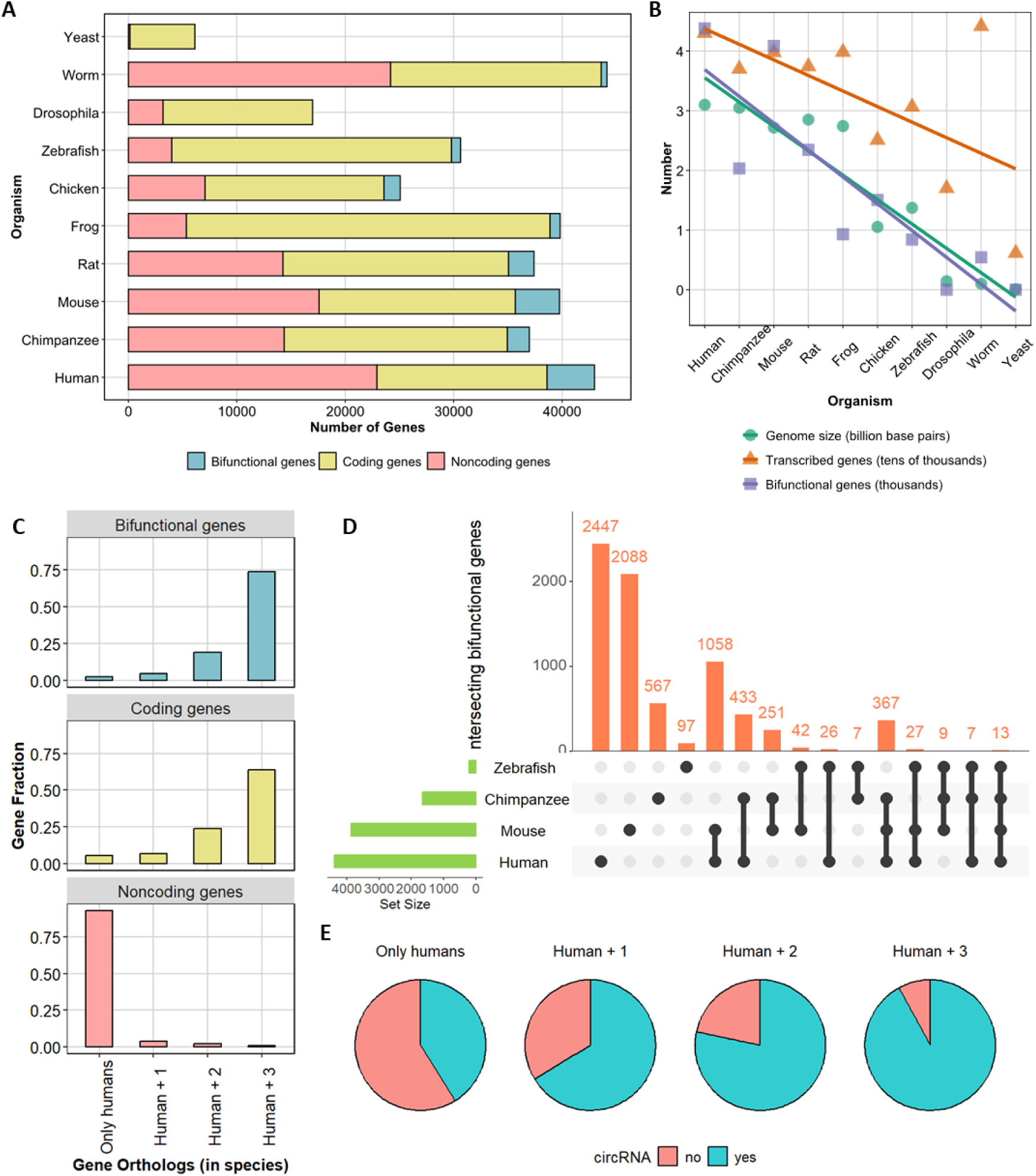
Bifunctional genes and evolutionary conservation. **(A)** The graph displays the distribution of bifunctional, coding and non-coding gene numbers across the genomes of ten model organisms. **(B)** The scatter plot show the correlation between Is there a relationship between genome sizes, number of bifunctional genes and the total number of transcribed genes across genomic time-lines from yeast to human genome **(C)** Presence of functionally conserved bifunctional genes across four organisms (Human, mouse, chimpanzee and zebrafish), we checked whether human bifunctional genes have orthologs in one, two or all of three other key model organisms. **(D)** Human bifunctional genes versus those in chimpanzee, mouse and zebrafish. Genes labelled as ‘bifunctional’ in any of the four organisms were combined to a bifunctional gene superset. This superset was then used to plot the number of overlapping bifunctional genes in all possible organism combinations (each bar represents the number of unique genes in the highlighted set). **(E)** Pie charts representing the number of bifunctional genes with annotated circular RNAs in each set: the ‘Only humans’ set represents genes with no mapped orthologs in Chimpanzee, Mouse, and Zebrafish. The ‘Human + 1’, ‘Human + 2’ and ‘Human+3’ represent the sets of bifunctional genes with orthologs in one, two or all three species respectively.

The observation that the genome of *Saccharomyces cerevisiae* (budding yeast), the most studied model organism does not harbour even a single bifunctional gene led us to examine transcript annotations of five more fungal genomes. Intriguingly, no bifunctional genes could be identified in any of these species which included well-studied multicellular fungi (Figure S3C). Non-coding genes constitute less than 2% of genes in five out of six fungal species analysed here. This could be an indication of minimal non-coding regulation in these organisms. Only the fission yeast, *Schizosaccharomyces pombe* is an exception, with 7,289 non-coding genes and yet with no bifunctional genes.

Sixty percent of plant genes are reported to undergo alternative splicing.^42^ Hence, we extended our analyses to include three well characterized plant species, namely *Triticum aestivum* (wheat), *Oryza sativa* (rice) and *Arabidopsis thaliana* (thale cress). The number of bifunctional genes varied widely among the three plant species, with Arabidopsis genome showing the least number of bifunctional genes (Figure S3D and Data S3). In general, the number of bifunctional genes in the three plant genomes were proportional to the number of non-coding genes, total transcribed genes as well as the genome size. However, since plant genomes are complex and often display polyploidy which has been shown to affect alternative splicing, it is not sure whether this may be extrapolated to all plants.^42–44^

### Bifunctional genes have conserved orthologs across species

To answer how bifunctionality is linked to conservation of genes and their functions across species, we went on to examine the following: first, to what extent are conserved genes bifunctional across the species (i.e. having orthologs) and the second, whether ‘bifunctionality’ of genes is conserved across species. To answer these questions, we mapped human genes to orthologs in three species (chimpanzee, mouse and zebrafish). Many transcribed genes in the human genome, specifically those annotated as ‘lncRNAs’ or ‘pseudogenes’, did not map to other species: only about 47% of human genes had orthologs in *Pan troglodytes* (chimpanzee). This number reduced to 45% and 40% in *Mus musculus* (mouse) and *Danio rerio* (zebrafish) respectively. Genes that mapped to all three species were considered ‘conserved’ with about 31% of all human genes falling into this category. This conserved fraction comprised of 73% of all bifunctional genes, 63% of all protein-coding genes, and less than 10% of all non-coding genes (Figure 2C). Thus, our findings suggest that compared to genes which are purely coding or non-coding, bifunctional genes show higher presence of orthologs across species.

Another interesting question was whether bifunctionality of genes (defined by the presence of coding and non-coding transcript variants) is conserved across species, which we answered by doing inter-species comparisons of bifunctional genes from four species. Only 13 genes are bifunctional across all 4 species analysed and 367 genes are bifunctional across the three vertebrate genomes studied (Figure 2D and Data S4). Out of the thirteen conserved genes *(CLASRP, HNRNPDL, KHDRBS1, KHDRBS2, NOP56, PLEKHA5, RPS9, SNX25, SRSF1, SRSF2, TASP1, UBP1, VCPKMT),* seven are involved in splicing/alternative splicing, whereas three others are RNA-binding proteins, showing a potential role for these conserved factors in posttranscriptional RNA processing.

Complex alternative splicing, sometimes arising from DNA variants, is the reason for bifunctionality and the diversity of transcripts from a gene can be further enhanced by back-splicing events, resulting in circular RNA transcripts. Over 20% of human genes are known to generate circRNAs at least in certain tissues.^45,46^ We mapped known human circular RNAs to bifunctional genes and found that 3,807 of 4,378 bifunctional genes have at least one circular RNA transcript. To map this finding to the conserved bifunctional gene orthologs across species, we extended our analyses to zebrafish, chimpanzee and mouse transcriptomes as in Figures 2C-D. Our analyses indicate that over 75 - 95% of bifunctional genes with at least two other orthologs give rise to at least one circular RNA transcript (Figure 2E).

### Bifunctional genes are predominantly involved in intracellular/vesicular transport, DNA binding and glycosylation

After establishing that a significant number of human bifunctional genes are conserved across species, we further explored the biological functions of bifunctional genes. For this, we performed a gene ontology (GO) analysis and found several interesting pathways and processes that were differentially overrepresented in the sets comprising of only bifunctional, only coding or only non-coding genes. We found that bifunctional genes are involved in a wide variety of processes and pathways and their expression is linked to DNA-binding and vesicular transport or recycling of nucleic acids and cellular cargo (Figure 3A, 3D and 3G) (Data S5, S6 and S7). Similar results were obtained when we used KEGG, MKEGG and WikiPathways genesets for analysis (Figure S4 A-H).

**Figure 3.**
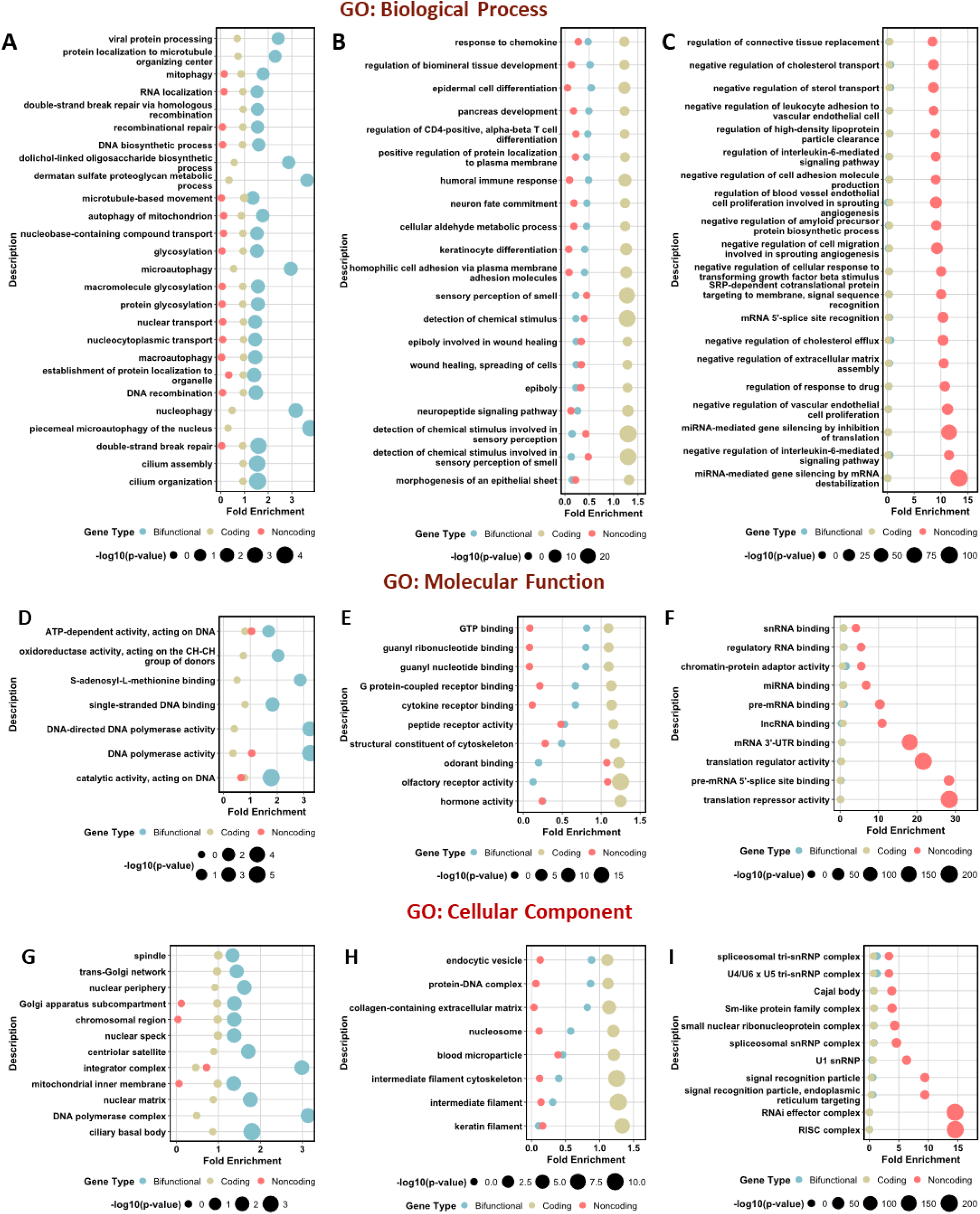
Bifunctional genes regulate fundamental biological processes distinct from coding and non-coding genes. Over-representation analysis of bifunctional genes (A,D,G) in comparison to coding (B,E,H) and noncoding genes (C,F,I) in various Gene Ontologies showing the top hits based on p-value cut-offs: GO: Biological Process (A-C), GO: Cellular Component (D-F) and GO: Molecular Function (G,H,I).

Since terms related to ‘DNA binding’, ‘DNA recombination’ and polymerase activity were enriched across all three families of GO, we examined the gene families which have more bifunctional members than others. Combining the ‘DNA recombination’ and ‘DNA biosynthetic process’ terms, we found that numerous DNA polymerase family members were bifunctional. This included the classical members *(POLB, POLL, POLM, POLI, POLK),* as well as the accessory and catalytic subunits (*POLD1, POLD4, POLE, POLE2, POLE3, POLG2)*. In addition to these, members of gene families from a multitude of processes were bifunctional, including lysine methyltransferases (*KMT5A, KMT5B*), helicases (*HELB, HELQM and HFM1*), several RNF family members (*RNF138, RNF212, RNF8*) and those associated with the mini-chromosome maintenance complex (*MCM2, MCM3 and MCM9)*.

Several members of the nucleoporin family (*NUP35, NUP43, NUP54, NUP83, NUP98, NUP160, NUP210*) as well as the SRSF family *(SRSF1, SRSF3, SRSF7*) which are members of the ‘RNA localization’ gene-set in in the Biological Process subcategory of GO are bifunctional. Furthermore, several autophagy-related genes (*ATG2A, ATG4A, ATG4D, ATG5, ATG7, ATG9A, ATG9B, ATG12, ATG13*) and sorting nexins (*SNX4, SNX6, SNX7, SNX14, SNX14, SNX18, SNX30*) also harbour coding and non-coding isoforms. Our analysis also revealed several terms related to glycosylation as being over-represented. More than 70 out of 200+ genes annotated in these terms were bifunctional with the ALG and GALNT gene families most enriched with 9 (*ALG1, ALG2, ALG3, ALG5, ALG8, ALG9, ALG11, ALG13, ALG14)* and 6 *(GALNT2, GALNT5, GALNTL5, GALNT11, GALNT13, GALNT14*, and *GALNT16*) bifunctional genes respectively.

We also categorized bifunctional genes based on the relative number of coding and non-coding variants annotated to them (Figure S2E). Interestingly, 350 bifunctional genes have more non-coding transcripts than coding transcripts. This subset of bifunctional genes also included genes involved in trafficking and autophagy (*RAB9B, RNASEK, ATG9B, ATG12*), genes from the SRSF (Serine/Arginine-rich splicing factor) as well as the TMEM (transmembrane proteins) and the ZNF (zinc finger proteins) families (*SRSF2, SRSF8, TMEM134, TMEM177, TMEM183A, TMEM222, TMEM67, ZNF154, ZNF431, ZNF514, ZNF525, ZNF594, ZNF689, ZNF705G, ZNF714, ZNF773, ZNF776*). The enrichment of these protein families in the bifunctional gene set with predominant presence of non-coding transcripts may indirectly indicate a role for associated non-coding transcriptome in the regulation of specific biological processes related to nucleic acid binding, RNA processing and vesicular transport.

To investigate whether the specific enrichment of certain GO terms with bifunctional gene sets are reliable, we performed similar analyses with pure ‘coding’ or ‘non-coding’ gene sets. When looking at coding genes, several terms related to chemokine/chemical stimuli response as well as differentiation and migration were significantly over-represented in the Biological Process sub-category of GO. In the ‘Molecular Function’ and ‘Cellular Compartment’ sub-categories, terms such as ‘receptor binding’ and ‘intermediate/keratin filaments’ were the top hits, suggest that coding genes are heavily involved in receptor-mediated signalling and cytoskeletal functions (Figure 3B, 3E and 3H). For ‘non-coding genes’, the top terms were those related to RNA binding and regulation of gene expression and translation, which is expected as most non-coding RNAs are characterised in these contexts (Figure 3C, 3F and 3I). Thus, gene ontology analyses show distinct differences in potential biological functions between coding, non-coding and bifunctional gene sets.

### Non-coding transcript variants of bifunctional genes are expressed in a cell and tissue specific manner

The long non-coding transcriptome is usually expressed at much lower levels than the coding mRNAs, which was the reason why LncRNAs were neglected as transcriptional noise in the past.^47^ Certain studies also show that the non-coding transcript variants of protein coding genes are expressed at very low levels at steady state.^13^ This is based on the assumption that they are targets of non-sense mediated decay (NMD). However, only around 2,000 of the 7,000+ validated non-coding transcripts arising from bifunctional genes are marked as NMD substrates (NCBI Gene Database: https://www.ncbi.nlm.nih.gov/gene/, accessed on April 01, 2024). To understand how coding and non-coding splice variants are expressed, we examined publicly available long-read and short-read RNA-sequencing datasets. To specifically assess the relative expression of coding and non-coding transcripts from bifunctional gene loci, we annotated exon-exon junctions unique to annotated coding and non-coding transcript variants from these genes. Using the exon-exon junction quantification function of the GTEx project, we then quantified these unique junctions and generated unique coding and non-coding fractions (summarized in Figure 4B). The tissue-median of noncoding fractions was over 0.05 for over 2600 bifunctional genes (Figure 4A and Data S8) and 778 genes had >5% non-coding junction reads in over 25 tissues (Figure 4C). Interestingly, over a hundred bifunctional genes had median noncoding fraction > 0.5 in more than 30 of 31 broadly categorized GTEx tissues (Figure S5A) indicating that noncoding junctions from these bifunctional genes are expressed ubiquitously at levels equal or greater than coding-specific exon-exon junctions.

**Figure 4.**
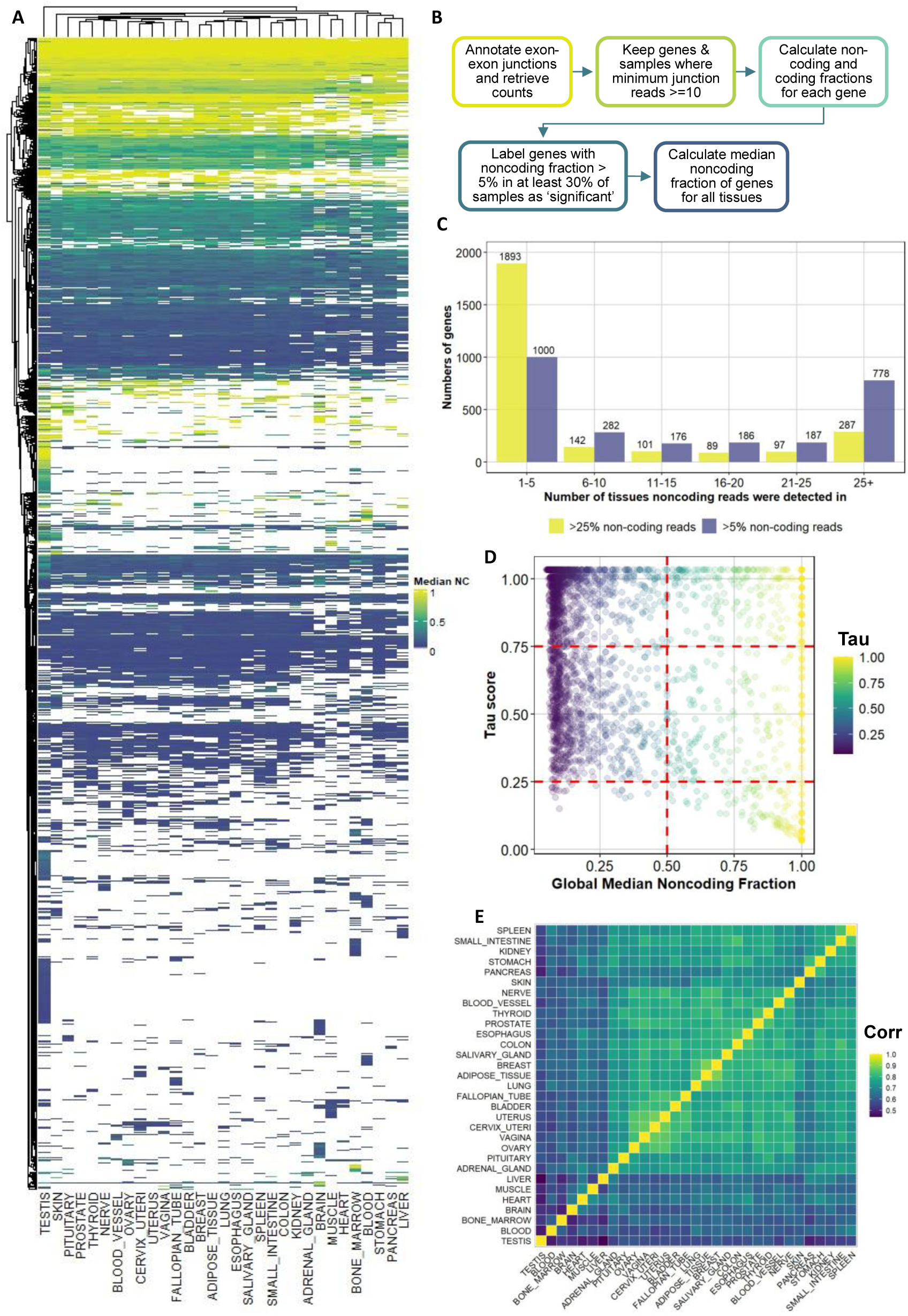
Expression of exon-exon junctions corresponding to noncoding transcripts of bifunctional genes in human tissues from GTEx. **(A)** Heatmap showing noncoding junction fraction was for 2609 bifunctional genes across 31 GTEx tissues, only genes which had >5% median noncoding fraction in more than 30% of the samples of a tissue are shown here. **(B)** Flowchart detailing the steps followed for exon-exon junction analysis. **(C)** Genes with non-coding junction expression across tissues with different cutoffs: green bars indicate number of genes with >5% noncoding junction reads in specified number of tissues whereas blue bars indicate number of genes with >25% noncoding junction reads. **(D)** Tau specificity score versus global median noncoding fraction for 2609 bifunctional genes. **(E)** Correlation between median noncoding fraction of 2609 bifunctional across different tissues in GTEx.

To understand the tissue specificity of noncoding isoform expression, we calculated Tau scores from median tissue expression for all 2600+ bifunctional genes showing non-coding variant expression based on unique non-coding junction quantification. Contrasting these tau values with global noncoding fraction medians for bifunctional genes show both categories of bifunctional genes - those with ubiquitous expression or with tissue specific expression of non-coding variants (Figure 4D). Furthermore, the expression of noncoding-isoform specific junctions seems to be highly correlated within tissues of epithelial origin and more tissue-specific in brain, muscle, blood and testis (Figure 4E and Data S9). Ten out of the thirteen conserved bifunctional genes had detectable noncoding fraction (>0.05) in at least one of 31 tissues (Figure S5B).

To further validate the expression of non-coding variants from bifunctional genes, we looked in detail for exons unique to their non-coding RNA splice forms. Out of the 13 conserved bifunctional genes, 7 genes (*CLASRP, HNRNPDL, KHDRBS1, NOP56, RPS9, SRSF1, and SRSF2*) harboured unique exon regions specific to non-coding variants, which were then probed through longread RNA-sequencing data available as part of the SGNEx (Singapore Nanopore Expression) project (Figure 5).^49^ Non-coding exon specific signals were detected in at least two of the six lines for all genes tested and consistent with the data for non-coding genes, the expression of the non-coding exons was mostly lower than coding exons of the same gene (Figures 5B-H). Interestingly, in case of the gene *CLASRP*, the non-coding transcript specific extended exon was detected across all cell lines (Figure 5A). The serine/arginine-rice splicing factor family members, SRSF1 and SRSF2, also displayed non-coding isoform expression in at least three of the 6 cell lines. Previous studies have also shown that such splicing patterns are characteristic of the SRSF gene family and are involved in internal splicing control.^15^ In addition to the conserved genes, *FUBP1,* another transcriptional regulator, also gives rise to non-coding transcripts that are expressed in five out of six cell lines (Figure 5F). An in-depth analysis of the SGNEx data also verified the expression of thousands of non-coding transcripts from 2248 bifunctional genes and the noncoding variants constituted the dominantly expressed transcripts from 434 genes in at least one cell line (Figure S6, Table S7, Data S11 and Data S12). Using a similar approach, we also investigated whether exon-specific expression in Adult GTEx (Adult Genotype Tissue Expression) project tracks available on UCSC Genome Browser, which revealed varying degrees of tissue specificity in the expression of the non-coding exons (Figure S7).^50^

**Figure 5.**
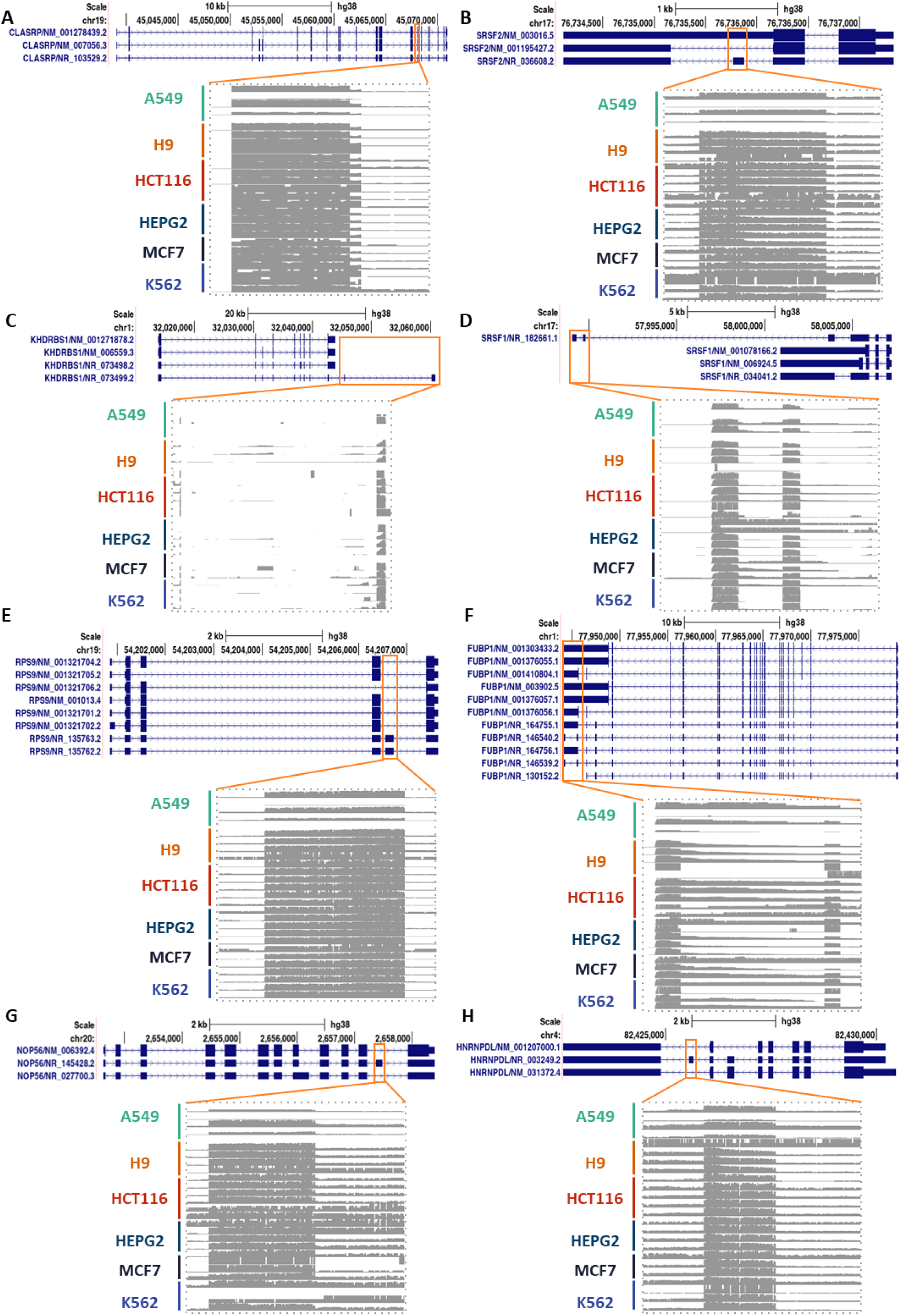
Monitoring expression of ncRNA variants by sequence reads from unique non-coding exons. Exon organization, showing the sequencing coverage of select conserved bifunctional genes from direct cDNA-based ONT RNA-seq data available from SGNEx. The reads for exon regions unique for non-coding transcripts are shown which indicate expression across 6 cell lines**. (A)** CLASRP has a 4-nucleotide extension in the highlighted exon which is unique to its noncoding isoforms. Unique non-coding exon expression for **(B)** *SRSF2*, **(C)** *KHDRBS1*, **(D)** *SRSF1*, **(E)** *RPS9*, **(F)** *FUBP1* (where the non-coding transcripts consist of 2 exons instead of a longer 3’-UTR), **(G)** *NOP56*, **(H)** *HNRNPDL*.

To experimentally verify the expression of these non-coding splice variants from the bifunctional genes, we designed non-coding RNA-specific primers (Figure S10) for select bifunctional genes. We used these primers to detect the expression of ncRNA isoforms from each gene by RT-qPCR in a panel of cell lines of diverse origin and found that these transcripts are expressed to detectable levels in many of the cell lines tested (Figure 5 and Table S5). More importantly, there exists cell-type specific expression patterns for ncRNA transcripts in some of these genes. For example, non-coding transcripts of the *DDX3Y* gene are expressed uniquely in only A549 (lung adenocarcinoma) and THP1 (monocytic cells) (Figure 5A), whereas the ncRNAs from *SNX16* and *XIAP* are expressed at detectable levels across all cell lines. In addition, non-coding variants of *USP15, TGFBR1* and *ACVR1B* are expressed at significantly lower levels in PANC1 (pancreatic cancer) cells when compared to A549 cells.

In addition to lineage specific differences, we also monitored the change in expression of ncRNAs from select bifunctional genes upon THP1 macrophage differentiation. A characteristic downregulation of non-coding transcript expression for *FUBP1, TIA1* and *FOXP1* was observed upon induction of THP1 macrophage differentiation and LPS stimulation (Figure 5B and Table S6).

### Non-coding transcripts of bifunctional genes may give rise to novel non-canonical proteins

Several lncRNAs have recently been shown to harbour non-canonical small open-reading frames (smORFs), that encode short polypeptides termed as microproteins (miPs).^51^ One of the proposed roles of the non-coding transcript variants of protein-coding genes is mediated by miPs that interfere with the function of the protein encoded by the gene.^31^ Smaller proteoforms encoded by splice-variants interfering with the functions of the canonical protein isoform has already been reported in case of *ZBP1* gene.^34^ To investigate the scope of similar regulation in bifunctional genes (Figure 1C), we predicted the potential ORFs harboured by their non-coding transcripts. Since non-canonical start codon (non-AUG) translation has now been reported as a widespread phenomenon, we utilized all possible start codons for our predictions.^52–54^ With a minimum length cut-off of 150 nucleotides, we predicted 1,25,660 unique ORFs (details in Table 1 and Data S10), most of which if translated, will give rise to microproteins (Figures 7A & 7B). The start codons of these predicted ORFs also lie mostly in the first quarter of the transcript’s length (Figure 7C) and about 20% of all ORFs are present in more than one non-coding transcript. The presence of smORFs in these non-coding transcripts further corroborates our incomplete understanding of the microprotein landscape as more and more proteins/proteoforms smaller than the canonical (>100 amino acid) polypeptides continue to be discovered.^55,56^

**Table 1.**
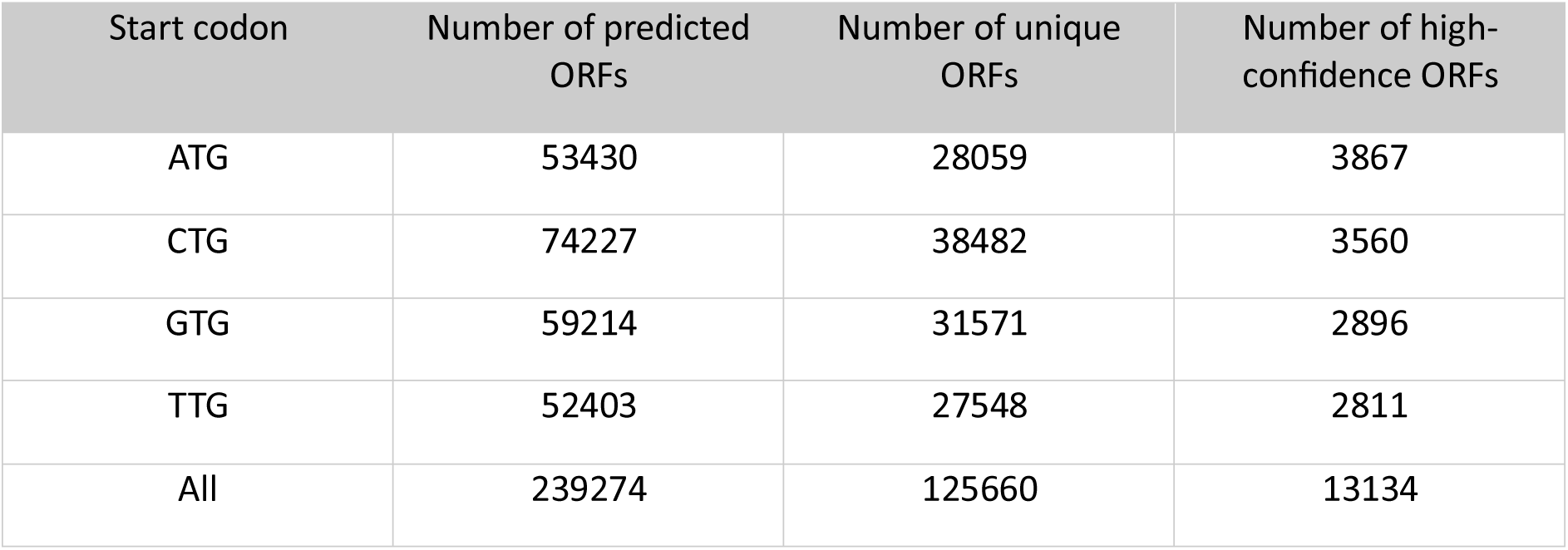
Summary of number of ORFs by start codon.

| Start codon | Number of predicted ORFs | Number of unique ORFs | Number of high-confidence ORFs |
| --- | --- | --- | --- |
| ATG | 53430 | 28059 | 3867 |
| CTG | 74227 | 38482 | 3560 |
| GTG | 59214 | 31571 | 2896 |
| TTG | 52403 | 27548 | 2811 |
| All | 239274 | 125660 | 13134 |

To further identify the truly translatable ORFs from our list, we calculated the translatability of these ORFs using multiple approaches. It should be noted that while our list includes only unique ORFs, avoiding redundancy from multiple ncRNAs harbouring the same ORFs, we do retain overlapping and nested ORFs to include all potentially translatable start-sites from a single transcript. Hydrophobicity of the C-terminus has been shown to influence the stability of translated peptides/proteins^57^ and Kozak context score gives a bioinformatic output indicating the translation potential of a particular start codon.^57,58^ We calculated the Kozak context score of the start codons and the C-terminal hydrophobicity of the encoded peptides to identify truly translated and stable miPs (Figure S9A). The subset of predicted ATG ORFs had a higher median Kozak context score than the ORFs with either CTG, GTG or TTG start codons (Figure 7D). Interestingly, GTG had the lowest median score, and the lowest number of ORFs with a score greater that the cut-off (>0.64) for higher translation probability. Even though the Kozak contexts different from one start codon to another, the hydrophobicity of the C-termini of the peptides arising from these ORFs did not differ between different start codon subgroups (Figure S9 B-C).

NMD is activated on mammalian mRNAs when the stop codon of the open reading frame is located upstream of an exon-exon junction.^59^ Incidentally, all major protein-coding ORFs in mammalian transcriptome have their canonical stop codons in the last exon. The presence of stop codons in the early exons is the reason why several transcripts with non-canonical ORFs are deemed substrates of NMD. To identify ncRNA variant-derived miPs with better stability, we blasted the predicted peptides to the accompanying annotated proteome and identified those that align with the C-terminus of known proteins. We then compared the Kozak context scores and C-terminal hydrophobicity of the peptides in three different groups namely (i) C-terminal blasting, (ii) non-C-terminal blasting and (iii) non blasting. As expected, the C-terminus hydrophobicity of non-canonical peptides was higher for non-blasting peptides, indicating that blasting peptides are more stable at cellular levels (Figure 6E). Coincidentally, the Kozak scores were also higher for blasting peptides, even though translatability predictions based on start codon context is not directly linked to blast similarity (Figure 6D).

**Figure 6.**
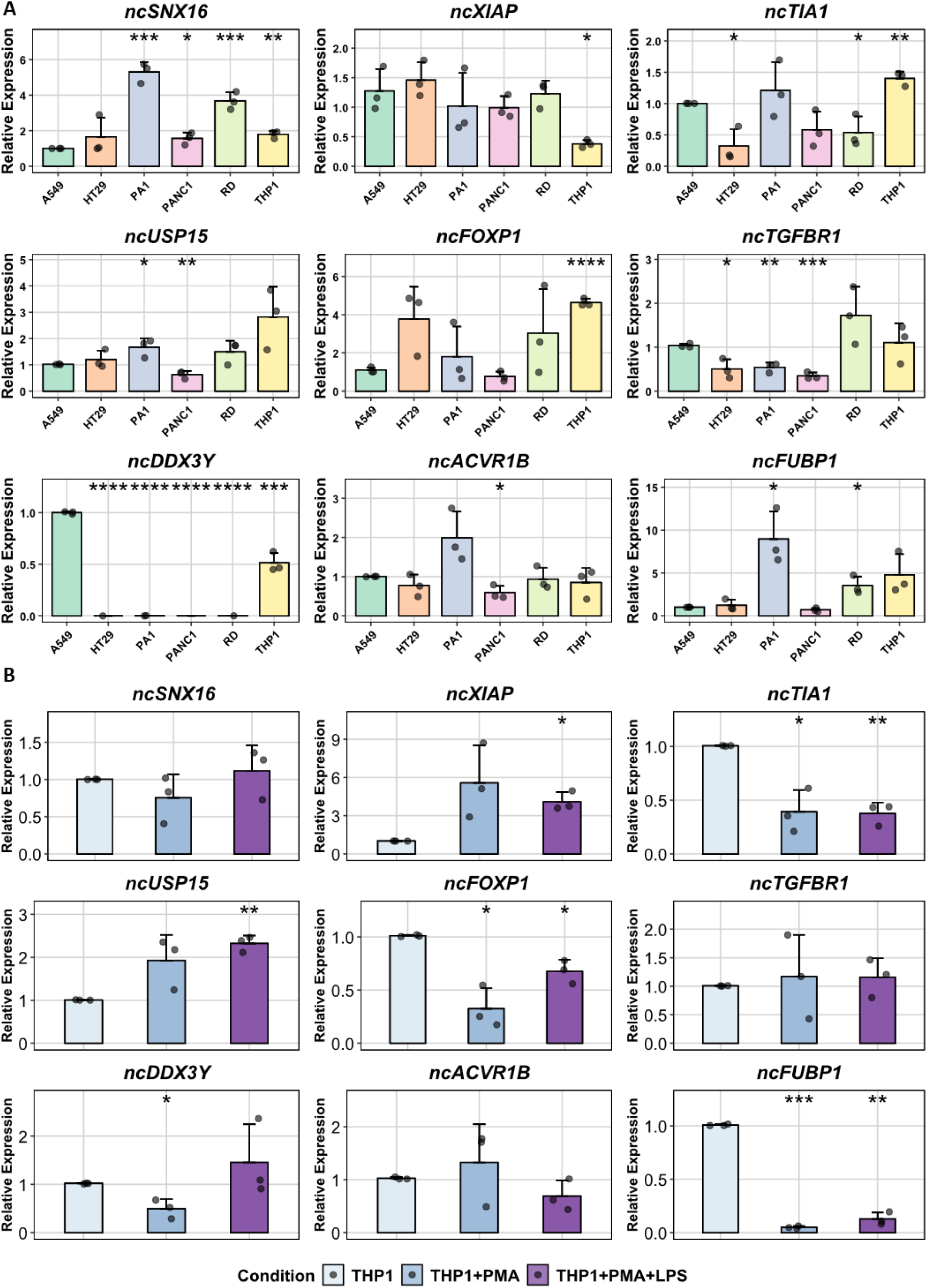
Cell line specific and stimulus specific expression of ncRNA variants from bifunctional genes. A &. **B.** RT-qPCR quantification of the expression of non-coding transcript variants from indicated genes were performed from a panel of cell lines to determine cell-type specificity **(A)** or in control or PMA differentiated THP1 monocytic cells (**B**). The PMA-differentiated THP1 macrophages were also immune primed further with lipopolysaccharide (LPS) treatment. p-values are indicated (n=3): *≤0.05; **≤0.01; ***≤0.001; **** ≤0.0001).

**Figure 7.**
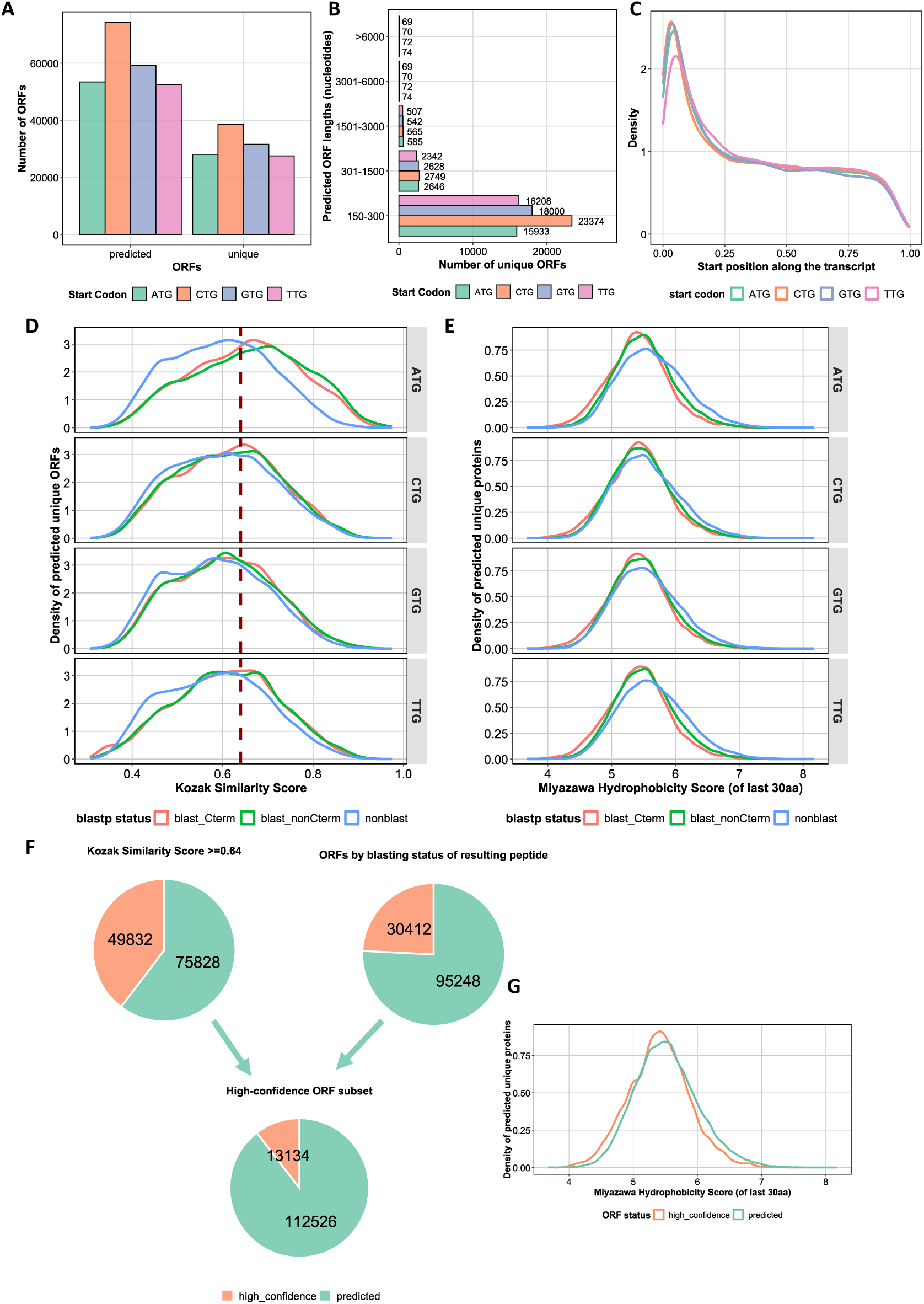
Are the non-coding transcripts of bifunctional genes truly non-coding? Translatability and stability of predicted proteoforms. **(A)** All predicted ORFs versus unique ORFs from non-coding transcripts for different start codons. **(B)** Size versus lengths of unique ORFs which may be translated into novel proteo-forms. **(C)** Start codon position of the predicted ORF relative to the complete transcript. **(D)** Kozak Score distribution of unique ORFs for start codons in combination with their *blastp* status. The dashed line marks the Kozak score of 0.64, above which the translation probability of ORFs is higher/accurately predicted for both cognate and non-cognate codons. **(E)** Miyazawa score of the last 30 amino acids indicating the C-terminal hydrophobicity of proteins arising from unique ORFs for different start codons in contrast with their *blastp* status. **(F)** Defining a subset of high-confidence unique ORFs based on Kozak Similarity Score (>=0.64) and *blastp* status of the resulting peptides. **(G)** Comparison of C-termini hydrophobicity of peptides from high-confidence ORFs versus other predicted ORFs.

While these ORF predictions are good start points for analyses, experimental verification is necessary to assess the true coding potential of these ORFs. To enhance the credibility of the predicted ORFs and to generate a group of high-confidence noncanonical ORFs, which are more likely to be translated and interfere with canonical protein function, we filtered the ORFs based on high Kozak Scores and C-terminal blasting status of resulting peptide. This subset consists of 13,134 ORFs which can be used for further experimental studies (Figure 6F & Figure S9E). Interestingly, we did not see a significant difference in the C-terminal hydrophobicity of the high-confidence subgroup (Figures S9 D & Figure 6G). Our predictions are further corroborated by independent predictions in the *OpenProt* server. Several of our predicted ORFs appear in *OpenProt* with supporting ribosome profiling and mass spectrometry evidence.^33,60^ Some of these predicted ORFs are expected to translate into shorter or slightly different proteoforms of the canonical protein product and may regulate their function as previously reported.^34^

### The “Bifunctional Genes Database” for exploring bifunctional gene architecture and potential regulatory functions

We summarized our findings in a publicly available and searchable ‘Bifunctional Genes Database’ available at https://janki-insan.shinyapps.io/bifunc_database/, which provides outputs from several analysis modules as presented in Figure 8. In addition to bifunctional gene information for humans and nine other species, the database also provides information on predicted ORFs and links to other resources for aiding comprehensive characterisation of individual bifunctional transcripts.

**Figure 8.**
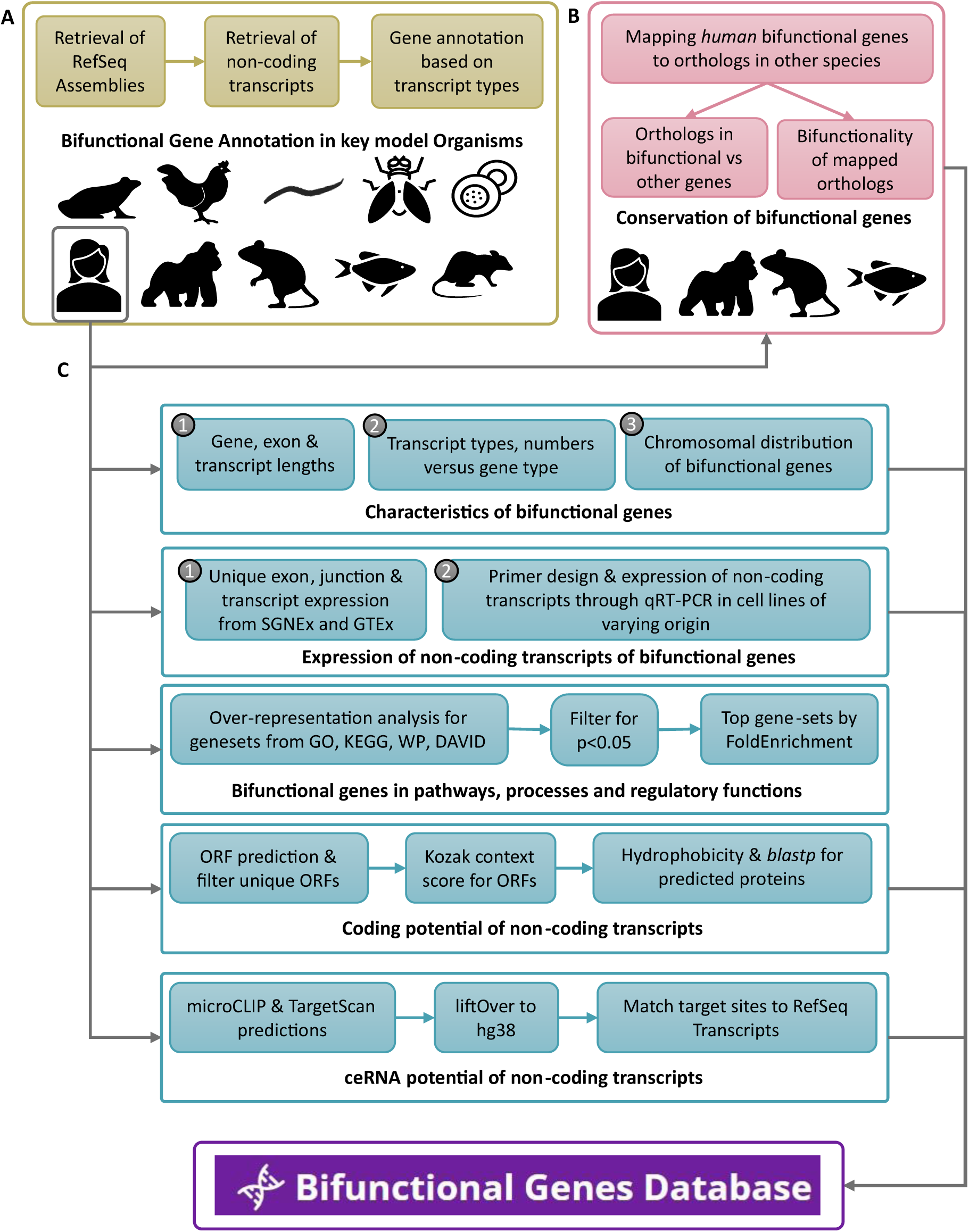
Summary of bifunctional gene analyses and features of the bifunctional genes database. **(A)** annotation of bifunctional genes, **(B)** their conservation across species and **(C)** understanding the influence of bifunctional genes on the human genome. The features of the searchable database are distributed into the above categories, and they are linked to information from other indicated servers for easy analyses. The database is available at https://janki-insan.shinyapps.io/bifunc_database/.

The non-coding transcript variants of protein coding genes may participate in the regulation of the coding transcript function as competing endogenous RNAs (ceRNAs) or miP encoding transcripts (^31,32^, Figure 1C). To determine the potential of non-coding transcripts as ceRNAs, we created a subset of overlapping high-confidence miRNA targeting seed sites from microCLIP and TargetScan and matched them with the transcript annotations.^61–63^ Over 100,000 miRNA-gene-transcript pairs were identified through both prediction approaches (Figure S12A and Data S14), which spanned over 100 human microRNAs and mapped to 3263 genes. In the common set, most human microRNAs targets about 15-20% of bifunctional genes, indicating that if expressed, non-coding transcripts could interfere with the silencing of their protein coding counterparts by microRNAs (Figure S12B). Within the bifunctional gene targets, we saw that for several genes, non-coding transcripts also contain the miRNA targeting seed sites (Figure S12C & S12D). Previous studies have reported this phenomenon^14^, however, further studies are required to investigate the mRNA/ncRNA expression ratios and the ability of the ncRNA variants to compete for miRNA binding. This potential function can also be explored with the help of the database.

## DISCUSSION

In our study, we performed a genome-wide analyses of human genes to reclassify them as coding, non-coding and bifunctional. We show that thousands of genes could potentially be ‘bifunctional’ and give rise to non-coding and coding transcripts regulated by alternative splicing events across organisms. The bifunctional genes are better conserved in evolution, and the number of bifunctional genes increase with genome complexity and size. Moreover, we also propose potential regulatory roles and miP encoding potential of these non-canonical transcripts. To facilitate further research on this special category of genes, we also generated a searchable database for bifunctional genes.

The non-coding transcriptome is often dismissed as transcriptional noise as lncRNAs are usually expressed at lower levels compared to mRNAs. A similar criticism can also be raised against the functional relevance of ncRNA variants from bifunctional genes. This may be supported by the possibility of many of them being NMD-targets and often expressed at low levels. However, research over the past decade have shown the functional relevance of several low expressed non-coding transcripts as regulatory RNAs. Thus, neglecting functional relevance of these transcripts due to low expression is not prudent. Moreover, there are comparable or a greater number of ncRNA variants expressed from several bifunctional genes, compared to mRNA variants. Previous reports have hypothesized the existence of one ‘dominant transcript’ and highlighted how non-coding splice isoforms rank as top expressed transcripts for 2119 human genes.^64^ This along with the exploration of whether dominant transcripts change in events of cellular stress, inflammation and hypoxia might be very interesting avenues to explore. The predicted ORFs and miRNA networks presented here will be useful tools for specific investigations into the regulatory mechanisms at each bifunctional gene locus. In addition to the mechanisms explored here, non-coding transcripts with appreciable expression levels, could compete with their mRNA counterparts for RNA-binding protein (RBP) binding, affecting posttranscriptional gene regulation at multiple levels. Distinctions are to be made between antisense transcripts and coding/non-coding variants generated from bifunctional genes, discussed here. While antisense transcripts could also be co-regulated or regulate the expression of its cognate sense-strand transcript, they are usually separate genes overlapping in opposite strands.

An interesting subset of the 4300+ bifunctional genes annotated here are associated with mRNA processing and RNA splicing. There is consistent over-representation of SRSF (serine/arginine-rich slicing factor) family genes in all our analyses. This is interesting as differential expression of SRSF genes can determine alternative splicing patterns. Expression of ‘poison-exon’ containing ncRNA transcripts from these genes has already been reported previously, especially in differentiation and tumorigenesis.^15,65^ SRSF1 and SRSF2 are among the 13 conserved bifunctional genes identified in our study and 7 of those genes *(CLASRP, HNRNPDL, KHDRBS1, KHDRBS2, SRSF1, SRSF2 and UBP1)* are directly involved in splicing. Moreover, alternative splicing has been shown to regulate several of these genes. *RPS9* orthologs in the *Saccharomyces cerevisiae* (budding yeast) and *Drosophila melanogaster* (fruit fly) are regulated by complex alternate splicing and intron inclusion.^66,67^ The *NOP56* gene is involved in ribosome biogenesis and snoRNA *SNORD86* derived from its intron regulates the alternative splicing and expression of the NOP56 protein, by controlling intron retention.^68,69^ When the entire bifunctional gene list is considered, we observe enrichment of other biological functions/processes which also include DNA binding, in addition to vesicular transport and autophagy. Interestingly, these involve fundamental processes key for survival and distinct from the overrepresentations shown for the purely coding and non-coding gene sets.

While we can see interesting evolutionary trends, the differences in depth of gene and transcript variant annotations across genomes should be considered while extrapolating these observations on genome complexity and the number of bifunctional genes. In addition to looking at the evolutionary conservation and functional enrichment of bifunctional genes, we also provide evidence of expression for non-coding transcripts of some of these bifunctional genes. While panning for unique ncRNA specific exons has been used to detect the presence of some ncRNA variants in long-read sequencing datasets, we also performed RT-qPCR analyses for experimental verification of ncRNA expression across a panel of cell lines of diverse origin which include A549 (lung), PANC1 (pancreas), THP1 (monocyte), HT29 (colon), PA1 (ovary) and RD (muscle) cells. Among the candidate bifunctional genes tested, *XIAP* (X-linked Inhibitor of Apoptosis) has been characterized as regulator of caspase-dependent cell death and has roles in immune disorders and cancer resistance.^70,71^ *TIA1* (T-Cell Intracellular Antigen 1) is a ubiquitously expressed gene which has been widely characterized for its roles in RNA splicing, cell cycle regulation, apoptosis and neuronal pathologies.^72,73^ Moreover, the expression of several non-coding transcripts for all the evolutionarily conserved and experiment validated genes discussed in the study has also been captured in CHESS (Comprehensive Human Expressed SequenceS V3.1.3) database, (Figure S4).^74^ Assessing the roles of non-coding transcripts in a cell- or tissue-specific manner for these bifunctional genes will uncover distinct layers of gene regulation.^72^ As an initial step towards this, we also looked at the expression of ncRNA variants of bifunctional genes during PMA-induced differentiation and pro-inflammatory priming of THP1 monocytes with lipopolysaccharide (LPS). *ncFUBP1* and *ncTIA1* were down regulated, and *ncXIAP* was up regulated during THP1 differentiation. Interestingly, these trends follow the expression patterns reported at the RNA level for these genes (Figure S11).^75^

Using the reference genome annotation for such analyses is just a start. The extent of how ncRNAs from bifunctional genes can influence its cognate mRNA expression and translation needs to be better understood in a context- or disease-specific manner. In that regard, there are studies which argue that non-coding isoform expression is only relevant when there is dysregulation of NMD, while others report widespread expression of non-coding isoforms, that may hint at the existence of more than 7,000 bifunctional genes.^13,76–78^ Most of all, as more long-read sequencing studies continue to discover novel isoforms, the number of bifunctional genes might continue to swell. As has been reported for currently understood bifunctional genes, analysis and annotations for such genes is required for all future studies to adequately address non-coding transcripts—whether they are expressed, are stable or they decay in a cell-type specific manner.^79^ Our analyses have shown that non-coding isoforms from ‘bifunctional genes’ represent a category of transcripts that cannot simply be overlooked. As has been previously reported, these non-coding transcripts might play very important roles in development and disease.^6,76,80^ Furthermore, exploration of key non-coding RNAs and accompanying genetic variants might give us newer clues of splicing-QTLs (Quantitative trait loci).^13^ In conclusion, the annotation of these genes as ‘bifunctional’ is necessary to facilitate research into their regulation.

### Limitations of the Study

As the first effort aimed at a genome wide analyses and reannotation of bifunctional genes, our work has several limitations. We have investigated and reannotated bifunctional genes based on the conservative RefSeq gene assembly (2023). While this was a necessary compromise to limit the analyses to better curated and validated transcript information, this would mean that the numbers we present here will evolve with emergence and validation of more and more long-read sequencing data in NCBI and GENCODE/ENSEMBL. While we have used isoform specific unique exons and unique exon-exon junctions to provide evidence for non-coding variant expression from proposed bifunctional genes, we are not able to assess expression from all bifunctional genes due to transcript and exon overlap. Several transcripts, even which have evidence of expression, might be candidates of non-sense mediated decay and/or without functional relevance. There are other cases, where non-coding transcripts might be reannotated as coding, after evidence emerge for their translation potential. The predictions of functional relevance, especially non-canonical translation needs to be supported by evidence from experimental studies. Furthermore, the microRNA predictions should only be used as a starting point of the analysis, not definitive evidence of possible function. While we have tried our level best to provide evidence for the expression of coding and non-coding variants from a vast majority of the human bifunctional genes, expression may not mean function. The term “bifunctional” is defined by the presence of potential coding and non-coding variants from a single gene and In-depth experimental evidence is required to prove dual functionality of the transcript variants.

## METHODS

### Sequence Retrieval and novel gene type annotation

The RefSeq transcriptome assemblies of the GRCh38.p14 (GCF_000001405.40-RS_2023_10) and T2T-CHM13v2.0 (GCF_009914755.1-RS_2023_10) genomes were retrieved from NCBI RefSeq database (https://www.ncbi.nlm.nih.gov/datasets/genome/) using the command-line tool datasets (v14.28.0)^96^. From the retrieved data, the RNA fasta file containing all annotated transcripts were traversed using Python (v3.10.13) and the number of coding and non-coding transcripts for each gene were counted. A comparison of both the RefSeq assemblies was done (results are in Table S3), and the GRCh38.p14 assembly (GCF_000001405.40-RS_2023_10), also referred to as ‘hg38’ was used for all further analyses. All transcribed genes in GRCh38.p14/hg38 were annotated based on whether they encoded only coding or non-coding transcripts, or both (labelled as ‘bifunctional’). For this annotation, all validated and predicted transcripts were considered (i.e. all transcripts accession IDs beginning with NM_, XM_, NR_ and XR_ were considered) and these labels were used for all further analysis. A further subset of the annotated genes was created using genes with only validated transcripts, which was used to check whether validated transcripts contribute to bifunctionality (see Figure S1).

### Bifunctional gene characteristics and Chromosomal location

To map genes to chromosomes and strands as well as compute other characteristics, the chromosomal location and strand information was retrieved from the genome annotation (.gtf) file accompanying the GRCh38.p14 assembly along with their current biotypes in NCBI. The gene fractions of bifunctional genes as well as their chromosomal distribution was plotted using *ggplot2* (v4.0.0) in R (v 4.4.2) (33,34). Genes present on both X and Y chromosomes were represented twice. i.e. on both the chromosomes and genes on alternate loci were represented as being on the parent chromosome (i.e. a gene at chr19_GL000209v2_alt was considered as being on and presented as chr19). The alluvial plots contrasting the NCBI biotype and our annotation was generated using the *ggalluvial* package(v0.12.5) ^82^. The characteristics of bifunctional genes versus others, such as gene lengths and number of transcripts were also computed using the genome annotation (.gtf) file. For number of exons, the number of exons in each annotated transcript of a gene was calculated and the transcript with maximum number of exons was used as representative of that gene. To compare bifunctional genes with their purely coding or non-coding counterparts, pairwise comparisons were made using the *stat_compare_means(method= ‘t.test’)* function from the *ggpubr* (v0.6.2) package. Effects sizes to compare the differences between the groups were also computed using the cohens_d() function from the *rstatix* (v0.7.3) package.

### Over-representation analysis of bifunctional genes

To determine the biological processes and pathways are more likely influenced by bifunctional’ genes, over-representation analyses of bifunctional genes, coding and non-coding genes was performed with genes of interest used as the test set and all annotated genes used as the background set. The analysis was performed for key GO genesets using the *enrichGO()* function and respective functions for KEGG, MKEGG and WikiPathways in *clusterProfiler* (v4.18.4) ^93,94^. For all function calls, the parameters pAdjustMethod = "BH”, pvalueCutoff = 0.01, and qvalueCutoff = 0.2 were used. Since the sub-ontologies ‘Biological Process’ and ‘Molecular Function’ list thousands of genesets, the top 25 genesets based on Fold Enrichment and p-values (p<0.05) were selected for plotting. Fold-enrichment and p-values for each gene-set from our results were plotted using *ggplot2* (v4.0.0) in R (v 4.4.2) ^97,98^.

### Conservation of bifunctional genes across species

To calculate the numbers of pure coding, non-coding and bifunctional genes in well-studied model organisms, the latest haploid RefSeq assemblies (accompanying the NCBI Reference Genome) for more than a dozen well-studied organisms across kingdoms were downloaded using the NCBI datasets tool and analysed using custom python scripts as described above (details in Table S3). Every gene harbouring a transcript was assigned a biotype. The transcript numbers for each organism (gene-wise) from this script were used to plot the total number of annotated genes, as done for humans, *ggplot2* (v4.0.0) in R (v 4.4.2) ^97,98^. Linear regression analysis to estimate the influence of genome size or the number of transcribed genes on the number of bifunctional genes was performed using the *lm()* function in R.

To check the interplay between conservation and bifunctionality in genes across related species, all human genes were converted to orthologs for three other species (chimpanzee, zebrafish, and mouse) using the *orthogene* (v1.16.0) package in R (v4.4.2) ^99^. Against the mapped orthologs, we compared gene fractions of each assigned gene type. In addition, we mapped genes that were bifunctional in other species and plotted them using *ComplexHeatmap* (v2.22.0) and *UpsetR* (v1.4.0) ^91,100^.

To map circular RNAs arising from bifunctional genes, we retrieved all human circular RNA transcripts annotated in the *circAtlas* (v3.0.0) and mapped them to bifunctional genes using their chromosomal coordinates in R ^30^.

### Expression of non-coding splice variants of bifunctional genes

To ascertain the expression of the non-coding variants of bifunctional genes, exon-exon junctions specific to coding, noncoding transcripts or shared between both transcript types were retrieved from the gtf file accompanying the October 2023 RefSeq hg38 release using the *GenomicFeatures* (v1.62.0)^95^, *tidyverse* (v2.0.0) and *txdbmaker* (v1.6.2) in R. These annotated junctions were matched to exon-exon junction expression data from GTEx tissues, retrieved from recount3 using the R package. From the mapped junction reads, reads shared between coding and noncoding transcripts of bifunctional genes were excluded and only genes and samples with at least 10 junction reads mapping to unique coding or noncoding junctions were retained. From the filtered data, the noncoding and coding fraction of junction reads was calculated as follows, i.e. Noncoding fraction = ∑(noncoding junction reads)/∑(coding junction reads + noncoding junction reads) and Coding fraction = ∑(coding junction reads)/∑(coding junction reads + noncoding junction reads). To denote a gene with sufficiently expressed noncoding junctions in a tissue, a cutoff of noncoding fraction > 0.05 (which corresponded to >5% noncoding junction reads) in more than 30% of the samples was placed. These filtered values were then used to calculate tissue-median noncoding fraction of remaining bifunctional genes. Global noncoding fraction medians and tau specificity scores (as previously described by Kryuchkova-Mostacci & Robinson-Rechavi^101^) were calculated using the above-defined tissue medians. To ensure the tau scores bifurcate between ubiquitously expressed and tissue-specific genes, a fixed denominator of n-1=30 (for 31 GTEx tissues) was taken.

To look at some genes in detail, aligned bigwig files from the Adult Genotype Tissue Expression (GTEx) Project and long-read RNA sequencing data for several cell lines available from the Singapore Nanopore Expression (SGNEx) project were visualized in the UCSC browser (https://genome.ucsc.edu/index.html).^2,49,102^ Expression of exons/junctions unique to non-coding transcripts in several bifunctional genes were visualized for bifunctional genes in the long-read data from SGNEx and short-read data from GTEx. In addition, fastq files from the direct-cDNA sequencing of various cell lines available as part of the SGNEx project were downloaded using AWS *s3 cli* (https://registry.opendata.aws/sgnex/). After quality inspection with NanoPlot, these files were mapped to RefSeq hg38 through minimap2 (v2.31) with the parameter *-ax splice:hq -uf -Y --MD*. Aligned bam files were sorted and indexed with samtools (v1.23.1) and input to isoQuant (v3.10.0) with the parameters *--data_type nanopore --report_novel_unspliced true --sqanti_ouput – polya_trimmed none* with the RefSeq fasta and gtf serving as reference and *genedb*. The ‘OUT.transcripts’ file with transcript-levels TPMs was used for all downstream analysis. Briefly, bifunctional genes with maximum TPM>=1 for at least one replicate were filtered out and the number of bifunctional genes expressing coding, noncoding or both kinds of transcripts were summarized. Next, a median TPM was calculated for each cell line was calculated for over 2000 matching bifunctional genes. In addition, the dominant transcript with the highest median TPM for each gene was ascertained for each cell line and plotted using *ggplot2* (v4.0.0).

Furthermore, the GTEx expression levels mapped to NCBI RefSeq transcripts curated as part of the CHESS3 (v3.1.3) database (https://ccb.jhu.edu/chess/) was also retrieved for selected genes using R and plotted using ggplot2 (v4.0.0) in RStudio.^50,74^ In addition, transcript annotations and accompanying TPM and sample count values were also retrieved from CHESS3 (v3.1.3).^74^ To plot the experimental RT-qPCR results (methodology described below), the packages ggplot2(v4.0.0), *cowplot* (v1.2.0), *ggpubr* (v0.6.2), *tidyr*(v1.3.1), *dplyr* (v1.1.4) and *reshape2* (v1.4.4) were used.^98^

### Primer design for non-coding splice variants

For select bifunctional genes, preferably intron-spanning primers specific to non-coding splice variants mapping either the exon skipping/intron retention or alternative splice site usage were designed using Primer3. The schemes for transcript-specific primer design of each gene are depicted in Figure S10. To ensure all primers were specific, they were blasted against the RefSeq RNA database and PCR amplification products resulting from each primer pair were run on agarose gels for size confirmation in addition to melt-curve analysis from RT-qPCR (data not shown).

### Cell culture, RNA isolation, cDNA synthesis and RT-qPCR analyses

A549 cells (ATCC, CCL-185) were a gift from Dr. Vamsi K. Yenamandra (CSIR-IGIB, New Delhi, India). RD cells were a gift from Dr. Sam J. Mathew (Regional Center for Biotechnology, Faridabad, India) and THP1 cells were kindly provided by Dr. Vivekanandan Perumal (IIT Delhi, New Delhi, India). All other cell lines were obtained from the National repository at NCCS, Pune, India. PA1, PANC1, HT29 and RD cells were cultured in DMEM High glucose media (Himedia, AL007A), A549 cells were maintained in DMEM/F12 (HiMedia, AL215A) and THP1 cells were maintained in RPMI (Himedia, AL028A). All media were supplemented with 10% fetal bovine serum (FBS) (Gibco, A5256701) and 1x antibiotic-antimycotic solution (Gibco, #15240062) and cells were grown at 37 °C, in a humidified atmosphere supplemented with 5% CO_2_. All cell lines were seeded in 6 well plates and grown for 24-48 hours till they achieved 80-90% confluency, following which cells were washed with PBS and lysed for RNA isolation. For THP1, 2 million cells were seeded in 6-well plates and differentiated using 100 ng/ml PMA (Cayman Chemicals, #10008014) for 24 hours, followed by a 24 hour recovery through media change. Differentiated THP1 cells were treated with 100 ng/ml LPS (Sigma, L2630) for 2 hours before the end of 24 hour recovery and used for RNA isolation.

RNA was isolated using the NucleoSpin RNA extraction kit (Macherey & Nagel) and 1ug RNA was used for cDNA synthesis using random hexamer primers according to the manufacturer’s instructions (Bio Bharati cDNA synthesis kit, BB-E0043). Real-time qPCRs for diluted cDNA (1:20) were run on a CFX96 (Bio-Rad) device using TB Green Premix Ex Taq II (Tli RNase H Plus, Takara) and the gene expression was normalized to Cyclophilin and relative expression calculated using the 2ΔΔCt method (46). All primers sequences are given in Table S4.

### Prediction of ORFs and their potential function

The non-coding transcripts (transcript IDs beginning with NR_ and XR_) arising from bifunctional genes were filtered from the original RNA fasta files using python (v3.10.13) in conda (v24.7.1). ORFs in these transcripts were predicted using ORFipy (v0.0.3) with the parameters *--min 150* (indicating minimum length of ORF) and *--strand f* (indicating the forward strand) for 4 possible canonical and non-canonical start codons (which included ATG, CTG, CTG and TTG) separately to ensure overlapping ORFs of importance are not missed.^81^ All predicted ORFs and resulting peptides were then given a unique ID based on the predicted sequence as more than one transcript could contain the same ORF. All predicted unique peptides were blasted against a proteome database created from all validated and predicted sequences from the GRCh38.p14 RefSeq release (GCF_000001405.40-RS_2023_10 contains 136,772 peptide sequences, 69,078 of which are predicted isoforms separately) using blast+(v2.14.0) with the following parameters, *-task blastp-fast -max_target_seqs 15 -outfmt 10*. A blast label was added to all predicted ORFs based on whether they blasted with the C-terminus of a validated/predicted protein to act as a flag for presence of premature termination codons, leading to non-sense meditated decay. To determine the translatability of the predicted ORFs, we calculated the Kozak context score previously described by *Gleason et al.*^58^ and the hydrophobicity scores of the resulting peptides from these ORFs were calculated using the peptides package (v2.4.6) in R (v4.4.2).^84^

### Predicting ceRNA potential of non-coding variants

To create a set of high-confidence non-coding transcripts from bifunctional genes that may compete for miRNA binding, chromosomal coordinates corresponding to miRNA seed sites retrieved from *microCLIP* and *TargetScan* were converted them to hg38 using the *LiftOver* executable with default parameters^61–63^. These chromosomal coordinates were then mapped chromosome wise to all transcripts (using the exon coordinates). Briefly, the exons closest to the microRNA seed sites was checked for overlap and overlapping sites from *microCLIP* and *TargetScan* with exons coordinates overlapping the seed sites and further analysis to determine the common targets from both studies in R (v4.4.2).

## Database

Output tables from above described analysis were used to create a reactive shinyApp using shiny (v1.12.1), with icons and graphics supported through bslib (v0.9.0) and bsicons (v0.1.2).^83,103,104^ The shinyApp is available at https://janki-insan.shinyapps.io/bifunc_database/.

## QUANTIFICATION AND STATISTICAL ANALYSIS

To quantify differences in transcript numbers, gene lengths and number of exons between bifunctional genes and other groups, the t-test from stat_compare_means in ggpubr (v0.6.2) was used. Effect sizes were also calculated between the groups using the effectsizes. All data for real-time qPCR is presented as mean ± S.E.M. Statistical significance on graphs is denoted by p ≤ 0.0001****, p ≤ 0.001***, p ≤ 0.01**, p ≤ 0.05*, p>0.05 (non-significant).

## RESOURCE AVAILABILITY

### Lead Contact

Further information and requests for resources should be directed to and will be fulfilled by the lead contact, Sonam Dhamija.

### Materials Availability

This study did not generate new reagents.

### Data and Code Availability

- **Data:** All datasets used in this study are publicly available. Processed & intermediate data is available in the Zenodo repository at www.doi.org/10.5281/zenodo.20478585. Source data for Figures is available in Tables S1- S7 or Data S1-S14. All datasets and annotations files used are publicly available from respective sources.
- **Database:** The shinyApp is available at https://janki-insan.shinyapps.io/bifunc_database/.
- **Code:** All analysis and figure scripts for this study are available at https://github.com/jankinsan/bifunctional_genes.

## Acknowledgements

S.D. thanks Department of Biotechnology (DBT), India for Ramalingaswami Re-entry Fellowship (BT/RLF/Re-entry/10/2019) for funding this work and DBT Wellcome Trust India Alliance for Intermediate Fellowship support (IA/I/22/2/506497). J.I. thanks Department of Education, Government of India for PhD fellowship. The authors thank Kusuma School of Biological Sciences and the HPC Facility, IIT Delhi for infrastructural support and computational resources. S.D. thanks CSIR-IGIB and South Asian University for infrastructural support.

## Author Contribution Statement

**JI:** conceptualization, methodology, investigation, analysis, software, writing, editing.

**MBM:** supervision, conceptualization, methodology, writing, editing.

**SD:** supervision, conceptualization, methodology, writing, editing, funding acquisition.

## Supplemental Information

**Document S1.** Figures S1–S12, Tables S1-S7, and supplemental references

**Data S1.** Gene characteristics for all genes from NCBI RefSeq GRCh38.p14 (GCF_000001405.40) (including the chromosomal and strand location), related to Figure 1

**Data S2.** Number of validated and predicted RefSeq transcripts from GRCh38.p14 (GCF_000001405.40) and their corresponding annotated gene type from our study, related to Figure 1 & S1

**Data S3.** A comparison of RefSeq assemblies across human assemblies and model organisms, related to Figure 2 & Figure S3

**Data S4.** Orthologs of human genes in mouse, chimpanzee and zebrafish along with their status in these organisms based on transcription products, related to Figure 2

**Data S5.** Over-representation analysis results for GO:BP through clusterProfiler, related to Figure 3

**Data S6.** Over-representation analysis results for GO:MF through clusterProfiler, related to Figure 3

**Data S7.** Over-representation analysis results for GO:CC through clusterProfiler, related to Figure 3

**Data S8.** Median non-coding read fraction and tau scores for 2609 bifunctional genes across GTEx tissues, related to Figure 4 & Figure S5

**Data S9.** Pearson correlation matrix for GTEx tissues based on median non-coding read fractions for bifunctional genes, related to Figure 4E

**Data S10.** Predicted unique ORFs from noncoding transcripts of bifunctional genes with ORF features and confidence labels, related to Figure 7 & Figure S9

**Data S11.** Ratio of TPMs of noncoding transcripts to the sum of TPMs for 1406 bifunctional genes, related to Figure S6D

**Data S12.** Dominant RefSeq transcripts of 2155 bifunctional genes with median TPMs across replicates for 7 cell lines profiled in SGNEx, related to Figure S6E

**Data S13.** Combined microCLIP and TargetScan predicted for all genes and human microRNAs, related to Figure S12

**Data S14.** MicroRNAs mapped to human bifunctional genes with transcript types, related to Figure S12

