## Supplementary material for "Genome-wide annotation and analyses of bifunctional genes in the human genome": Document S1

**Document S1.** Figures S1–S12, Tables S1-S7, and supplemental references

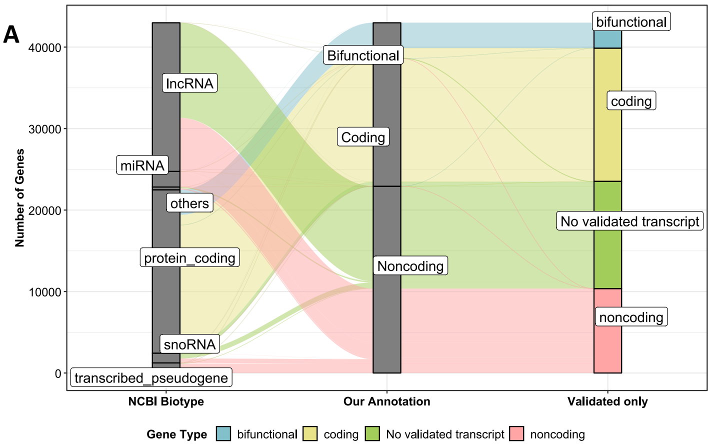


B

| Category | RefSeq Complete | RefSeq Validated |
|---|---|---|
| Transcribed genes | 42987 | 29820 |
| Bifunctional Genes | 4378 | 3114 |
| Coding Genes | 15689 | 16344 |
| Noncoding Genes | 22920 | 10362 |

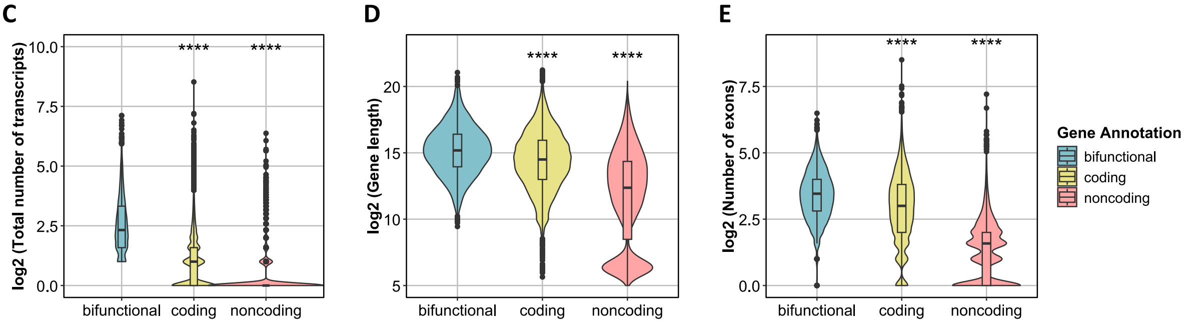


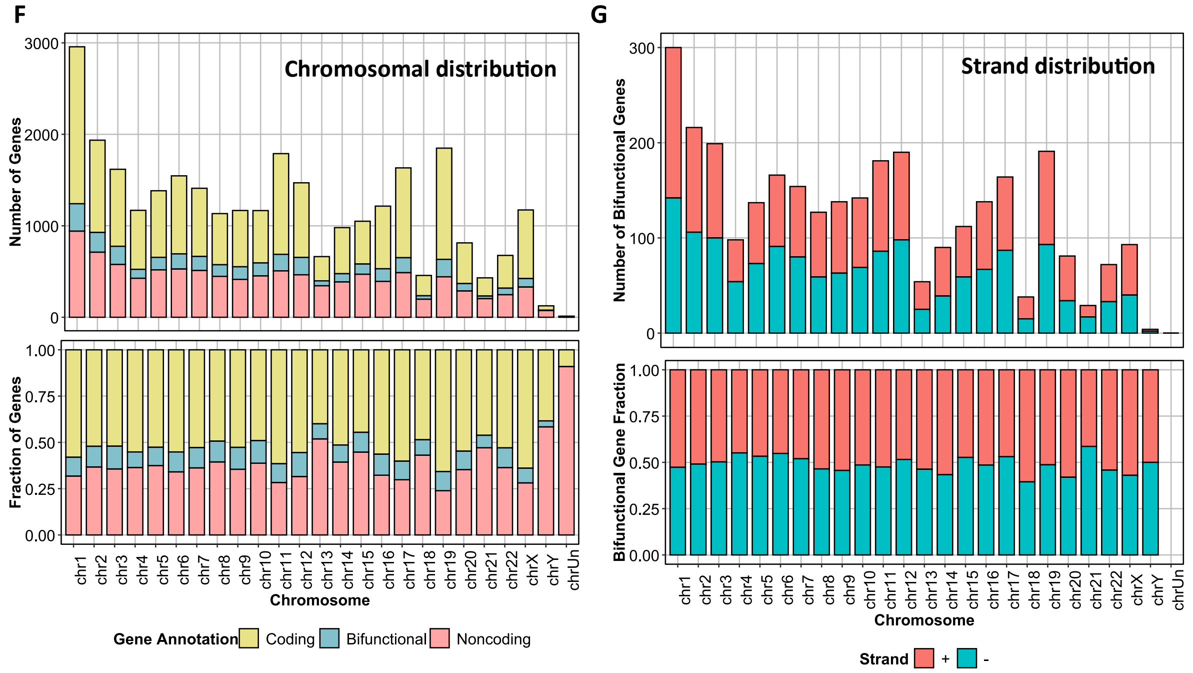


**Figure S1 Reannotation of bifunctional genes using RefSeq Validated transcripts. (A)** Alluvial plot showing the distribution of NCBI biotypes versus our annotation or reannotation with only validated transcripts. **(B)** Number of total transcribed, bifunctional, coding and noncoding based on Complete RefSeq versus validated RefSeq transcripts. (C-E) Differential analyses of number of transcripts (D), gene lengths (E) and number of exons in the longest transcript for bifunctional genes versus purely coding and non-coding genes for the genes with only validated transcripts (Pairwise comparisons were made using t-test, **** denotes p ≤ 0.0001, *** denotes p ≤ 0.001, ** denotes p ≤ 0.01 and * is used for p ≤ 0.05 respectively). Distribution of bifunctional genes with validated transcripts versus genes with validated mRNA and ncRNA transcripts on different chromosomes **(F)** and different strands **(G)**.

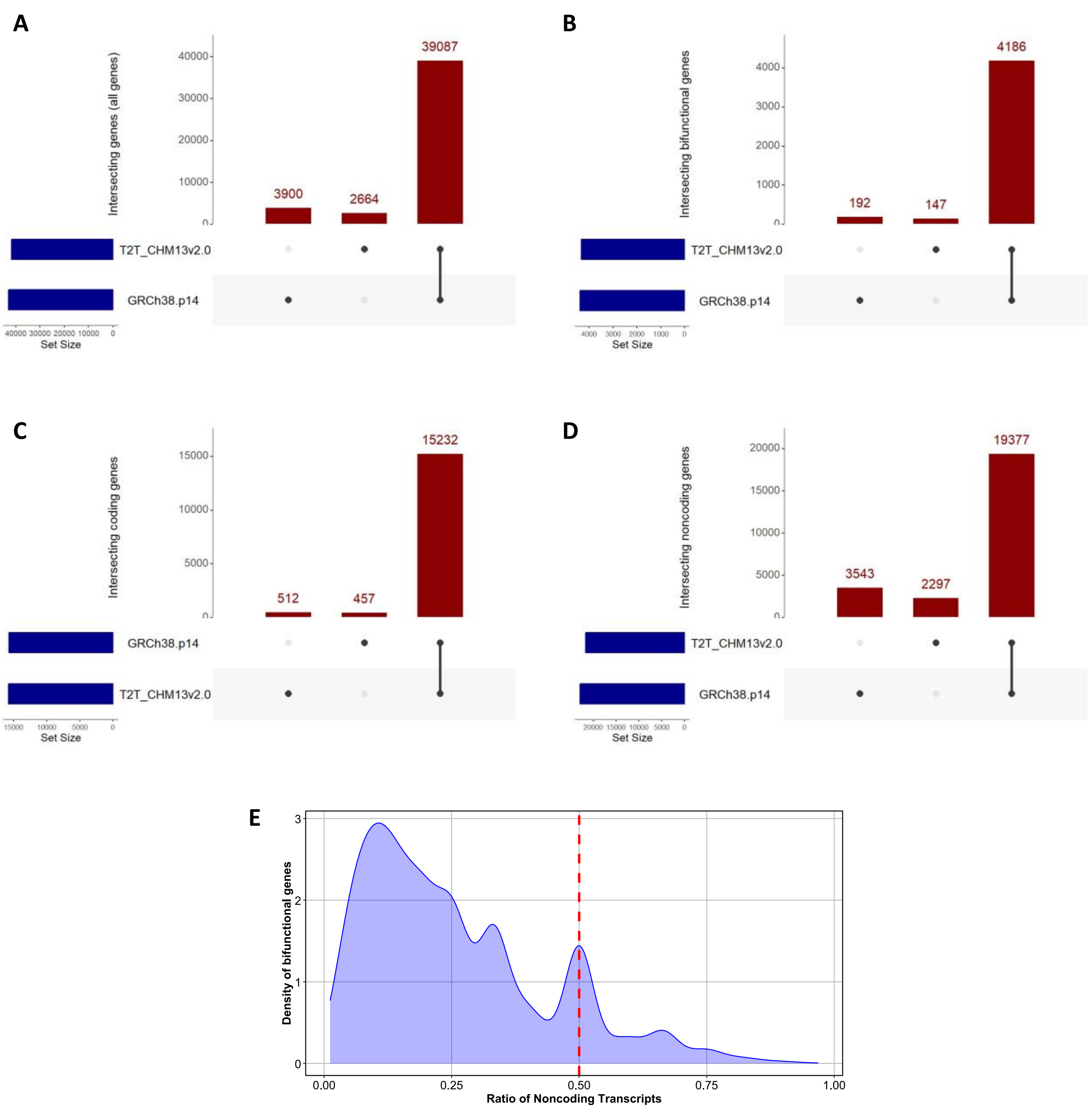


**Figure S2 (A), (B), (C)** and **(D)** are a comparison of all, bifunctional, coding and noncoding genes from the GRCh38 and T2T assemblies. **(E)** shows the density distribution of the ratio of annotated non-coding transcripts for bifunctional genes.

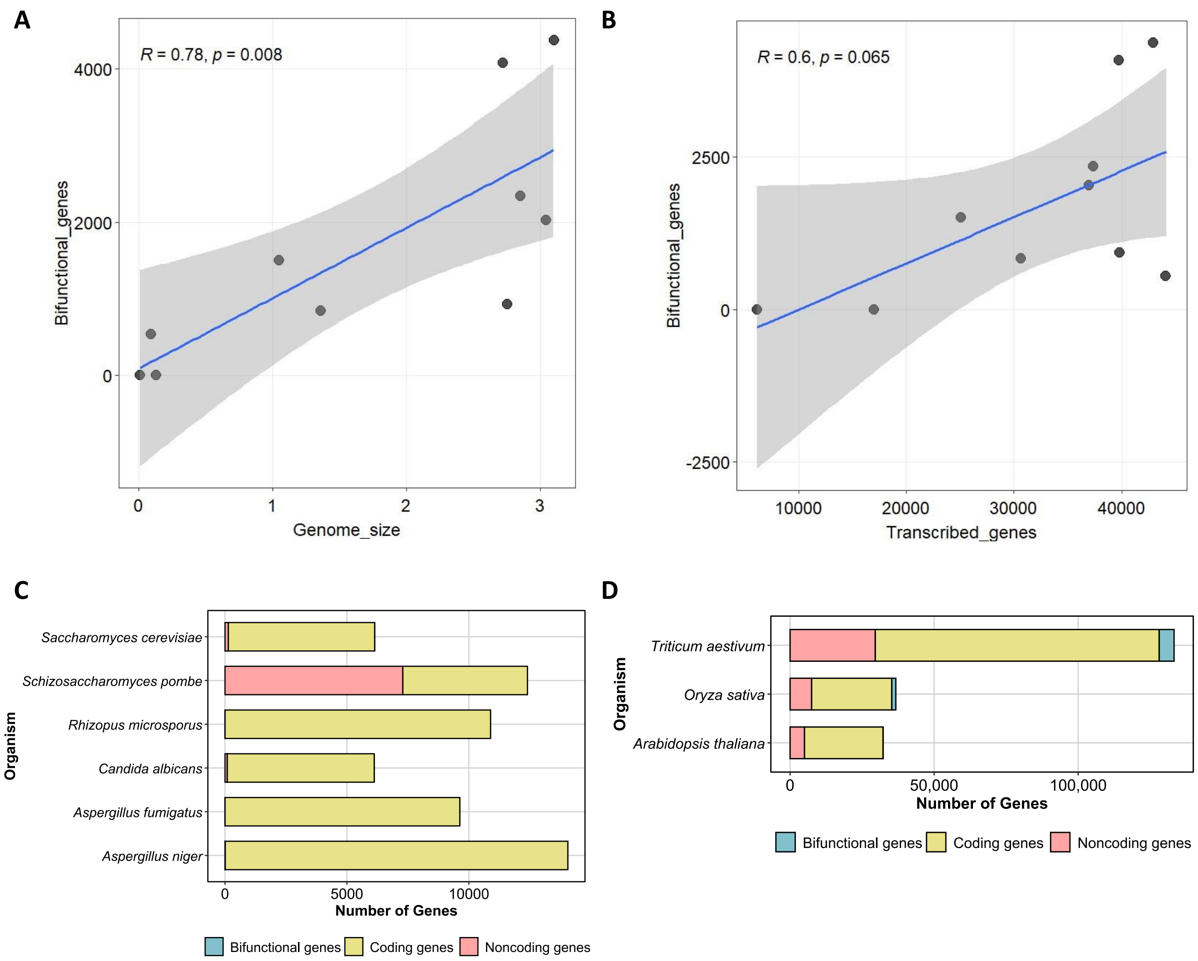


**Figure S3** Linear regression to predicted bifunctional gene number based on **(A)** Genome size, **(B)** Number of transcribed genes for key model organisms described in Figure 2. Comparison of bifunctional genes in fungal and plant species is presented in **(C)** and **(D).**

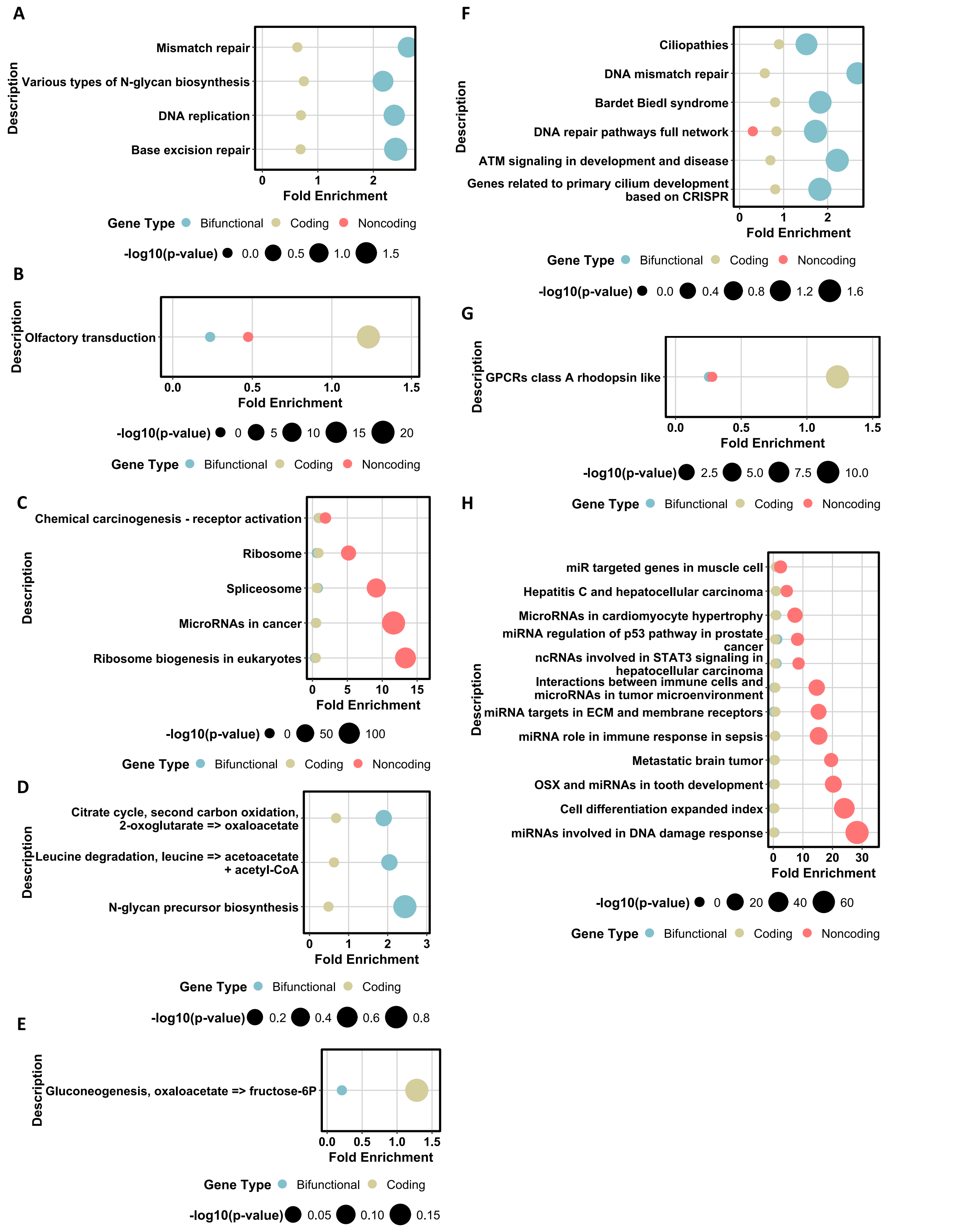


**Figure S4** Enrichment of bifunctional, coding and noncoding genes using genesets from KEGG **(A), (B) & (C),** MKEGG **(D), (E)** and WikiPathways **(F), (G) and (H).**

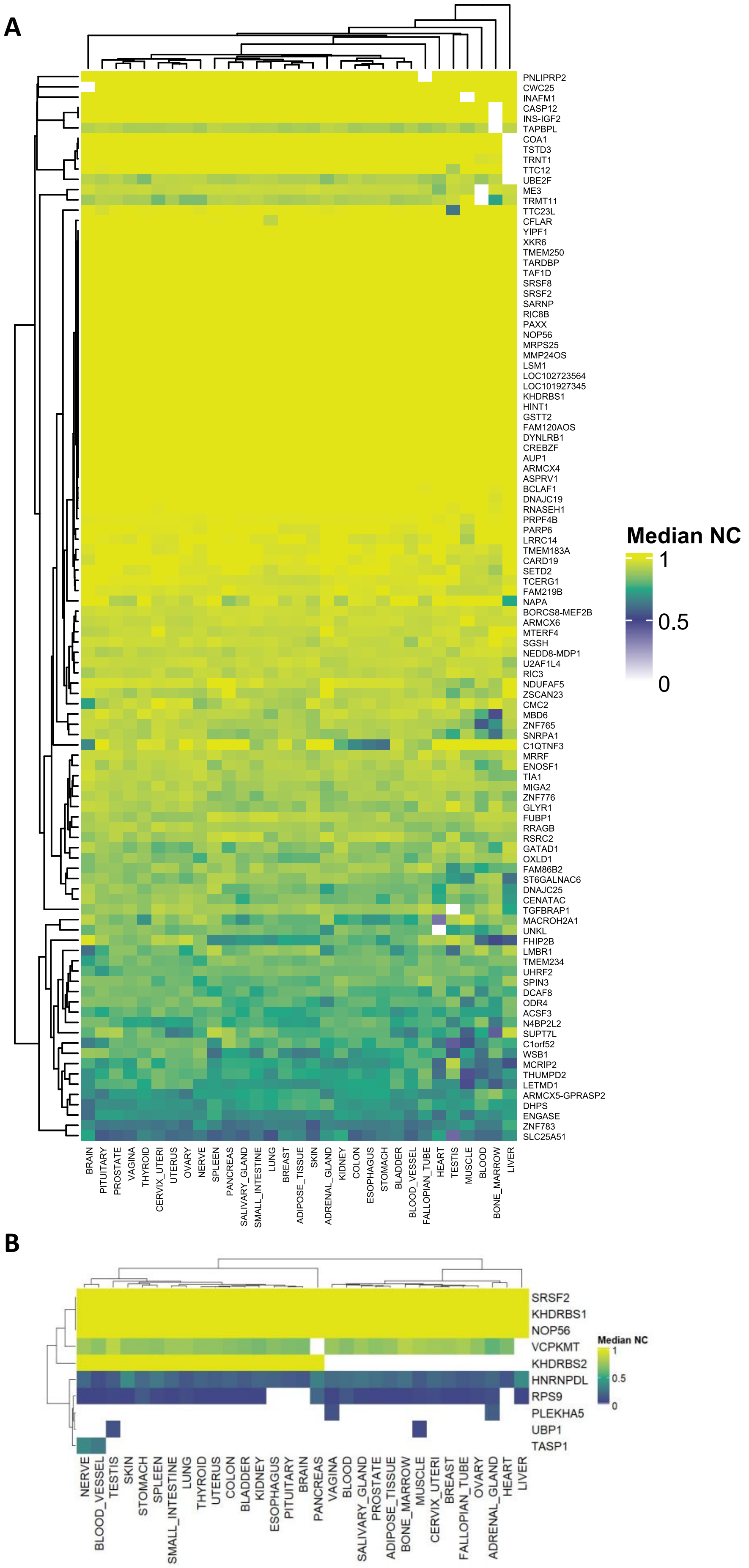


**Figure S5 (A)** Median noncoding fraction across tissues for 104 bifunctional genes with >50% noncoding reads in at least 30 of 31 GTEx tissues. **(B)** Median noncoding read fraction of conserved bifunctional genes across GTEx tissues (only 10 out of 13 genes where noncoding reads passed the cutoffs for detectable expression are plotted).

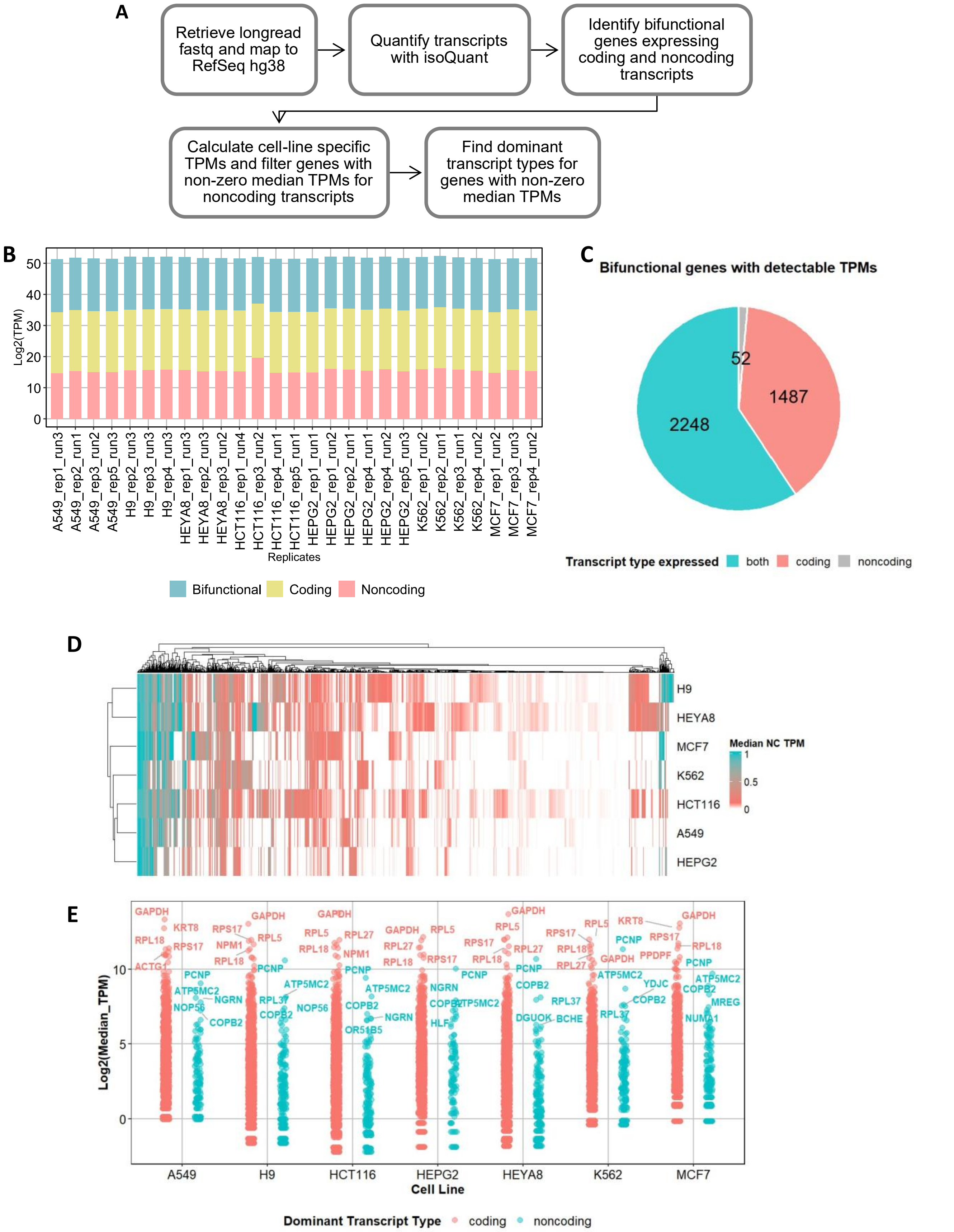


**Figure S6 Expression of coding and noncoding transcripts of bifunctional genes from SGNEx**[1] **(A)** Pipeline followed to quantify transcript expression from analysis of ONT sequencing of direct cDNA from various cell lines provided in SGNEx. **(B)** Sum of TPM values for annotated RefSeq transcripts across different cell lines and replicates. Since TPM sums for most replicates were close or above 900,000, novel transcripts were excluded from this analysis. **(C)** Transcript-type expression of bifunctional genes with the sum of TPMs was >=1 across cell lines and replicates. **(D)** Heatmap showing ratio of median TPM for noncoding transcripts to all transcripts for 1400+ bifunctional genes (represented by rows) across different cell lines (columns). **(E)** Dominant transcript type for bifunctional genes across cell lines. From the cell-line specific median TPM, a dominant transcript was denoted for each cell line and plotted. Top 5 genes with dominant coding and noncoding transcripts are labelled.

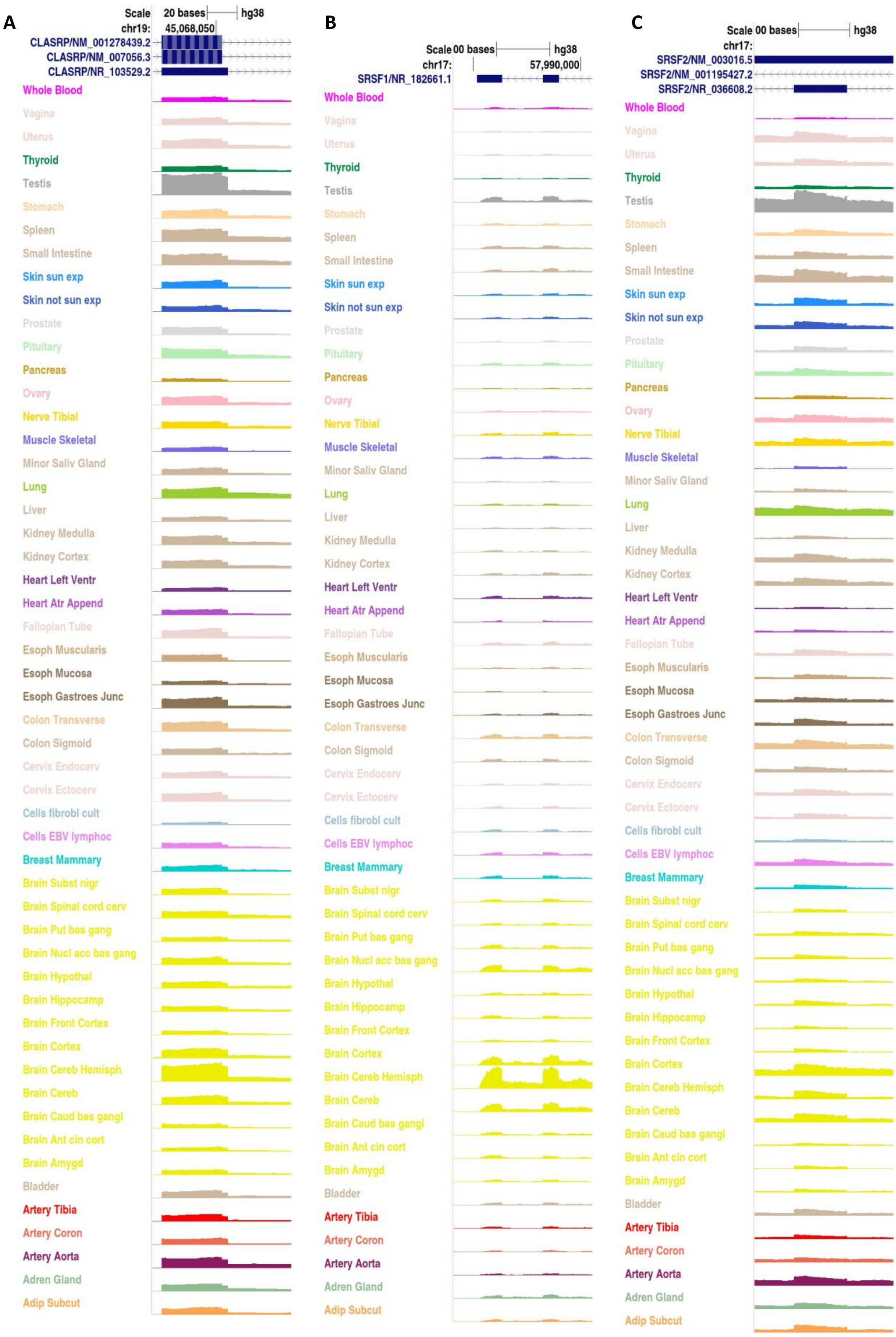
A
Scale 20 bases hg38
chr19: 45,068,050
CLASRP/NM_001278439.2
CLASRP/NM_007056.3
CLASRP/NR_103529.2
B
Scale 00 bases hg38
chr17: 57,990,000
SRSF1/NR_182661.1
C
Scale 00 bases hg38
chr17:
SRSF2/NM_003016.5
SRSF2/NM_001195427.2
SRSF2/NR_036608.2
Whole Blood
Vagina
Uterus
Thyroid
Testis
Stomach
Spleen
Small Intestine
Skin sun exp
Skin not sun exp
Prostate
Pituitary
Pancreas
Ovary
Nerve Tibial
Muscle Skeletal
Minor Saliv Gland
Lung
Liver
Kidney Medulla
Kidney Cortex
Heart Left Ventr
Heart Atr Append
Fallopian Tube
Esoph Muscularis
Esoph Mucosa
Esoph Gastroes Junc
Colon Transverse
Colon Sigmoid
Cervix Endocerv
Cervix Ectocerv
Cells fibrobl cult
Cells EBV lymphoc
Breast Mammary
Brain Subst nigr
Brain Spinal cord cerv
Brain Put bas gang
Brain Nucl acc bas gang
Brain Hypothal
Brain Hippocamp
Brain Front Cortex
Brain Cortex
Brain Cereb Hemisph
Brain Cereb
Brain Caud bas gangl
Brain Ant cin cort
Brain Amygd
Bladder
Artery Tibia
Artery Coron
Artery Aorta
Adren Gland
Adip Subcut

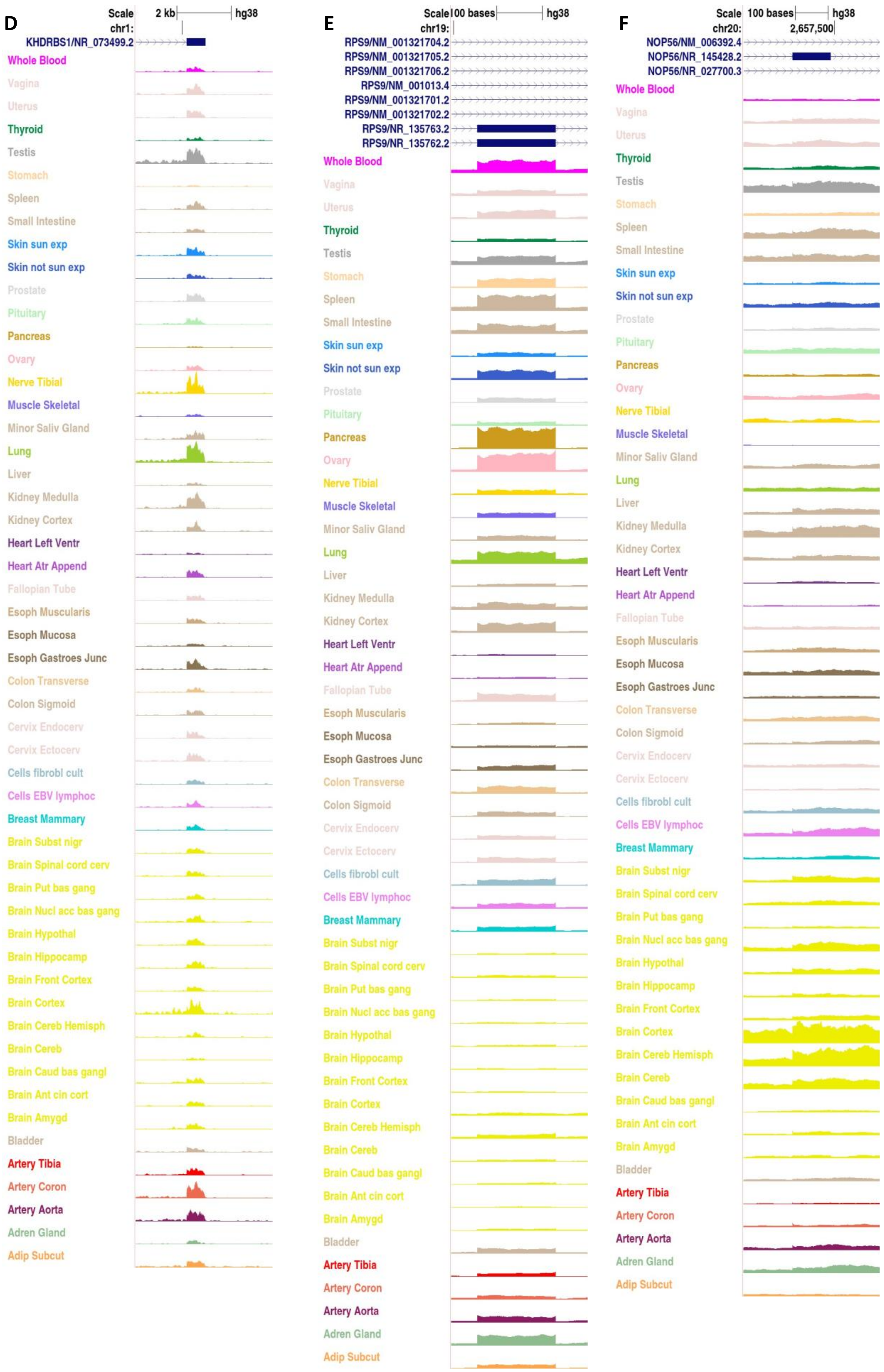
D
Scale 2 kb hg38
chr1:
KHDRBS1/NR_073499.2
Whole Blood
Vagina
Uterus
Thyroid
Testis
Stomach
Spleen
Small Intestine
Skin sun exp
Skin not sun exp
Prostate
Pituitary
Pancreas
Ovary
Nerve Tibial
Muscle Skeletal
Minor Saliv Gland
Lung
Liver
Kidney Medulla
Kidney Cortex
Heart Left Ventr
Heart Atr Append
Fallopian Tube
Esoph Muscularis
Esoph Mucosa
Esoph Gastroes Junc
Colon Transverse
Colon Sigmoid
Cervix Endocerv
Cervix Ectocerv
Cells fibrobl cult
Cells EBV lymphoc
Breast Mammary
Brain Subst nigr
Brain Spinal cord cerv
Brain Put bas gang
Brain Nucl acc bas gang
Brain Hypothal
Brain Hippocamp
Brain Front Cortex
Brain Cortex
Brain Cereb Hemisph
Brain Cereb
Brain Caud bas gangl
Brain Ant cin cort
Brain Amygd
Bladder
Artery Tibia
Artery Coron
Artery Aorta
Adren Gland
Adip Subcut
E
Scale 100 bases hg38
chr19:
RPS9/NM_001321704.2
RPS9/NM_001321705.2
RPS9/NM_001321706.2
RPS9/NM_001013.4
RPS9/NM_001321701.2
RPS9/NM_001321702.2
RPS9/NR_135763.2
RPS9/NR_135762.2
Whole Blood
Vagina
Uterus
Thyroid
Testis
Stomach
Spleen
Small Intestine
Skin sun exp
Skin not sun exp
Prostate
Pituitary
Pancreas
Ovary
Nerve Tibial
Muscle Skeletal
Minor Saliv Gland
Lung
Liver
Kidney Medulla
Kidney Cortex
Heart Left Ventr
Heart Atr Append
Fallopian Tube
Esoph Muscularis
Esoph Mucosa
Esoph Gastroes Junc
Colon Transverse
Colon Sigmoid
Cervix Endocerv
Cervix Ectocerv
Cells fibrobl cult
Cells EBV lymphoc
Breast Mammary
Brain Subst nigr
Brain Spinal cord cerv
Brain Put bas gang
Brain Nucl acc bas gang
Brain Hypothal
Brain Hippocamp
Brain Front Cortex
Brain Cortex
Brain Cereb Hemisph
Brain Cereb
Brain Caud bas gangl
Brain Ant cin cort
Brain Amygd
Bladder
Artery Tibia
Artery Coron
Artery Aorta
Adren Gland
Adip Subcut
F
Scale 100 bases hg38
chr20: 2,657,500
NOP56/NM_006392.4
NOP56/NR_145428.2
NOP56/NR_027700.3
Whole Blood
Vagina
Uterus
Thyroid
Testis
Stomach
Spleen
Small Intestine
Skin sun exp
Skin not sun exp
Prostate
Pituitary
Pancreas
Ovary
Nerve Tibial
Muscle Skeletal
Minor Saliv Gland
Lung
Liver
Kidney Medulla
Kidney Cortex
Heart Left Ventr
Heart Atr Append
Fallopian Tube
Esoph Muscularis
Esoph Mucosa
Esoph Gastroes Junc
Colon Transverse
Colon Sigmoid
Cervix Endocerv
Cervix Ectocerv
Cells fibrobl cult
Cells EBV lymphoc
Breast Mammary
Brain Subst nigr
Brain Spinal cord cerv
Brain Put bas gang
Brain Nucl acc bas gang
Brain Hypothal
Brain Hippocamp
Brain Front Cortex
Brain Cortex
Brain Cereb Hemisph
Brain Cereb
Brain Caud bas gangl
Brain Ant cin cort
Brain Amygd
Bladder
Artery Tibia
Artery Coron
Artery Aorta
Adren Gland
Adip Subcut

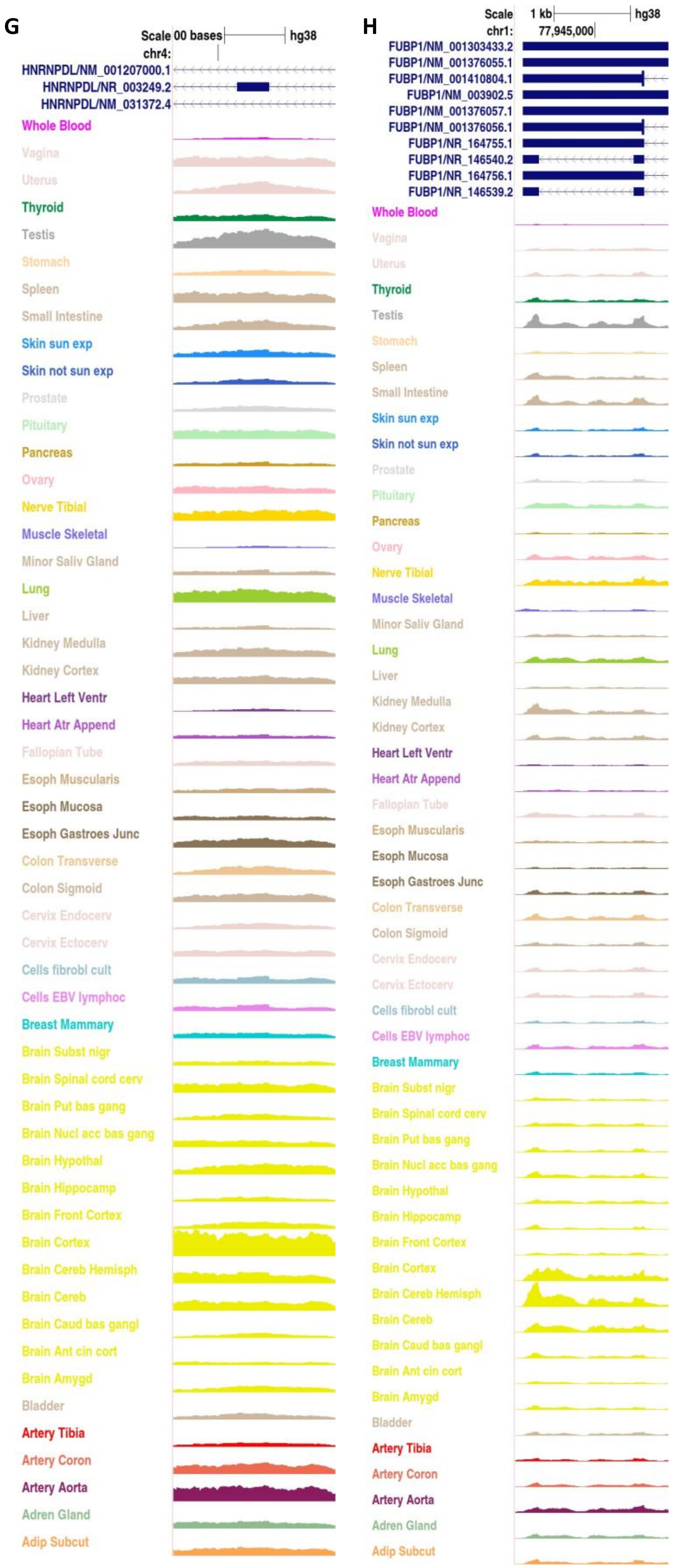


**Figure S7** shows the expression of noncoding isoform-specific-exon from GTEx tracks available in the UCSC Genome Browser (tracks available from https://genome.ucsc.edu/s/jankinsan/hg38_gtex_rna_seq_coverage).

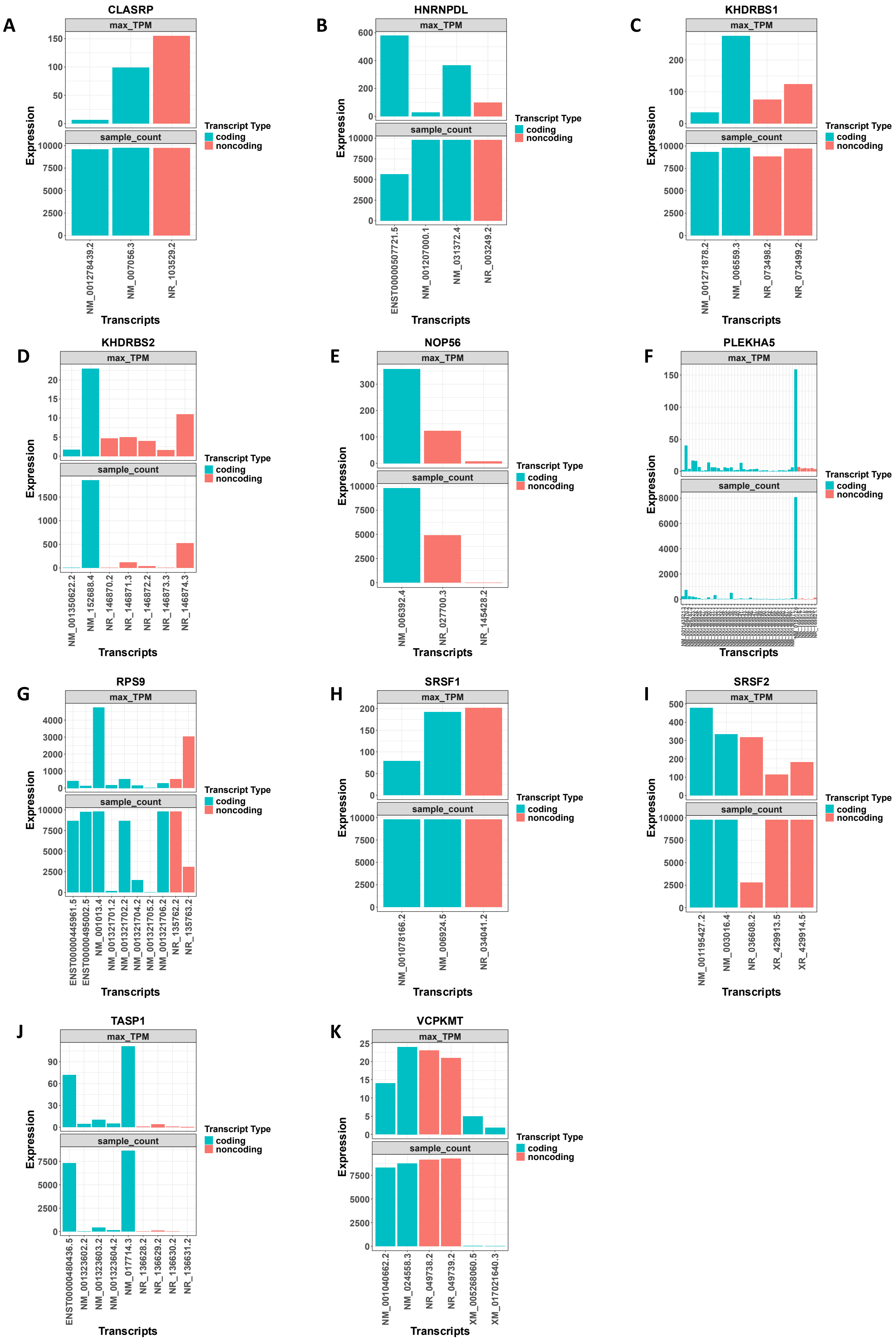
A
CLASRP
max_TPM
sample_count
Expression
NM_001278439.2
NM_007056.3
NR_103529.2
Transcripts
Transcript Type
coding
noncoding
B
HNRNPDL
max_TPM
sample_count
Expression
ENST00000507721.5
NM_001207000.1
NM_031372.4
NR_003249.2
Transcripts
Transcript Type
coding
noncoding
C
KHDRBS1
max_TPM
sample_count
Expression
NM_001271878.2
NM_006559.3
NR_073498.2
NR_073499.2
Transcripts
Transcript Type
coding
noncoding
D
KHDRBS2
max_TPM
sample_count
Expression
NM_001350622.2
NM_152688.4
NR_146870.2
NR_146871.3
NR_146872.2
NR_146873.3
NR_146874.3
Transcripts
Transcript Type
coding
noncoding
E
NOP56
max_TPM
sample_count
Expression
NM_006392.4
NR_027700.3
NR_145428.2
Transcripts
Transcript Type
coding
noncoding
F
PLEKHA5
max_TPM
sample_count
Expression
Transcripts
Transcript Type
coding
noncoding
G
RPS9
max_TPM
sample_count
Expression
ENST00000445961.5
ENST00000495002.5
NM_001013.4
NM_001321701.2
NM_001321702.2
NM_001321704.2
NM_001321705.2
NM_001321706.2
NR_135762.2
NR_135763.2
Transcripts
Transcript Type
coding
noncoding
H
SRSF1
max_TPM
sample_count
Expression
NM_001078166.2
NM_006924.5
NR_034041.2
Transcripts
Transcript Type
coding
noncoding
I
SRSF2
max_TPM
sample_count
Expression
NM_001195427.2
NM_003016.4
NR_036608.2
XR_429913.5
XR_429914.5
Transcripts
Transcript Type
coding
noncoding
J
TASP1
max_TPM
sample_count
Expression
ENST00000480436.5
NM_001323602.2
NM_001323603.2
NM_001323604.2
NM_017714.3
NR_136628.2
NR_136629.2
NR_136630.2
NR_136631.2
Transcripts
Transcript Type
coding
noncoding
K
VCPKMT
max_TPM
sample_count
Expression
NM_001040662.2
NM_024558.3
NR_049738.2
NR_049739.2
XM_005268060.5
XM_017021640.3
Transcripts
Transcript Type
coding
noncoding

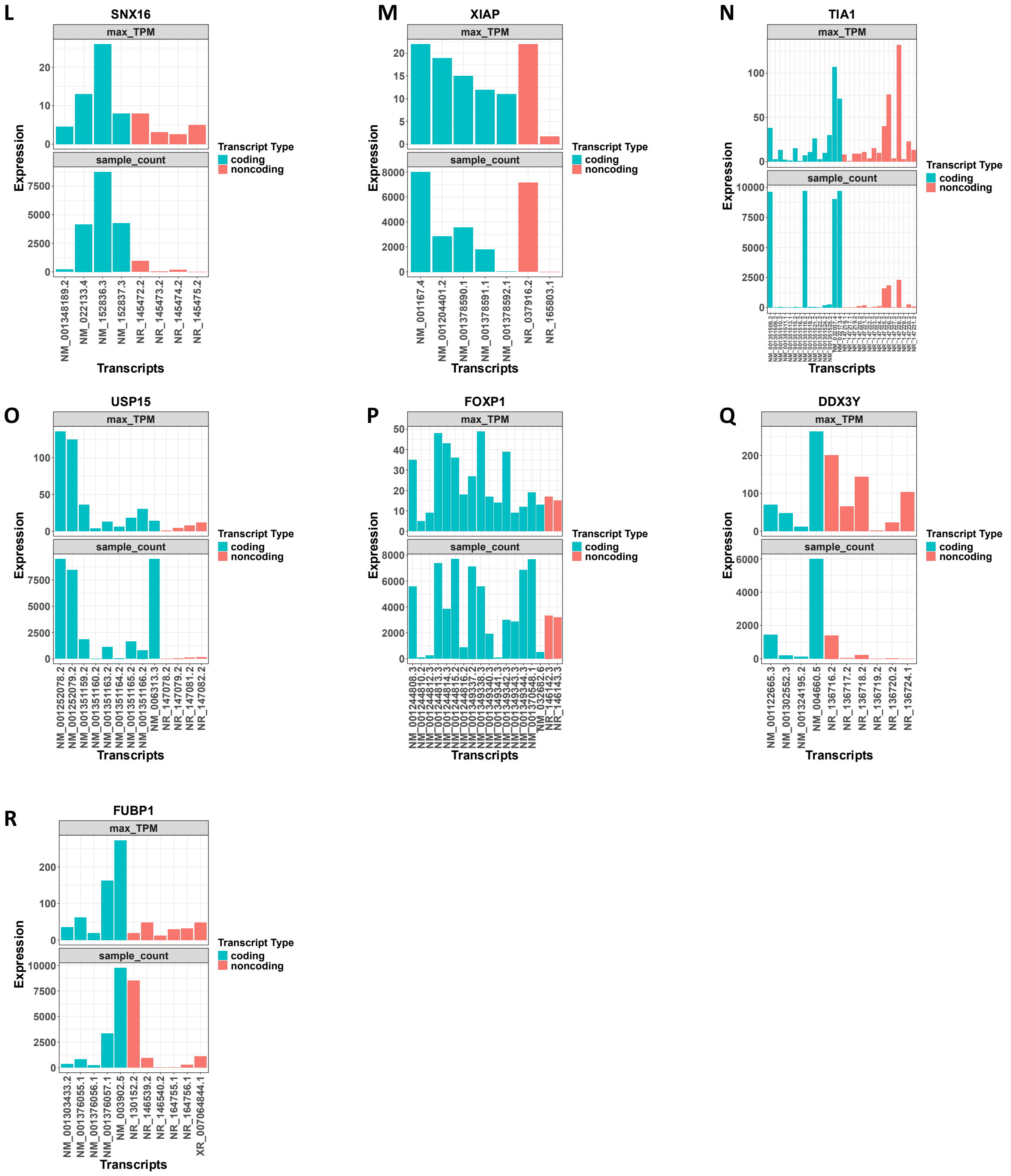


**Figure S8** shows the expression for non-coding transcripts from CHESS3[2] (https://ccb.jhu.edu/chess/). The maximum TPM and the number of samples the transcript was expressed as plotted as bar plots.

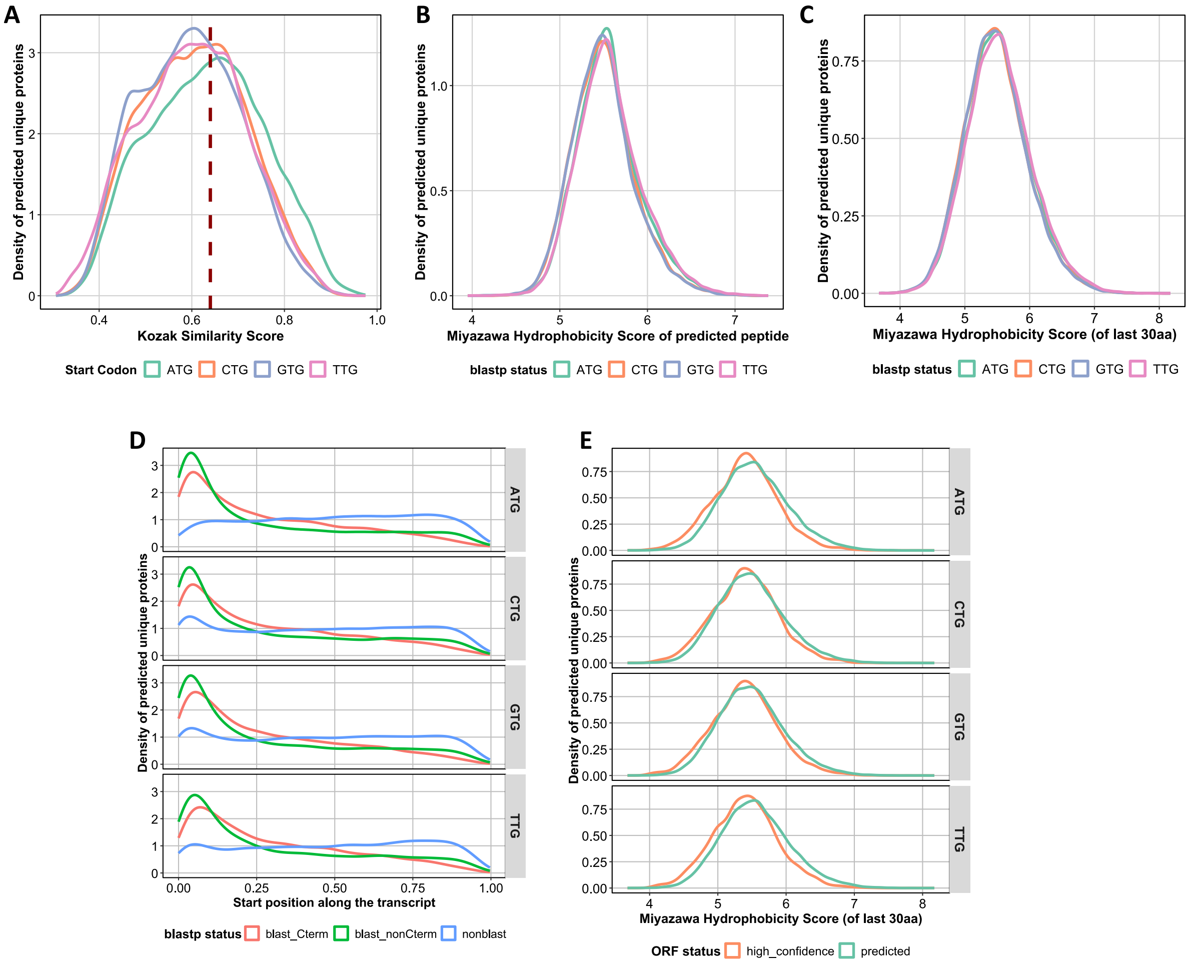


**Figure S9 (A)** Distribution of Kozak Score of predicted unique ORFs versus the start codon. The dashed line marks the Kozak score of 0.64. Distribution of Miyazawa Hydrophobicity scores for the predicted complete proteins **(B)** and the last 30 amino acids for different start codons **(C)**. **(D)** Distribution of relative start position for predicted ORFs with different start codons in contrast with the *blastp* status of the resulting peptides. **(E)** C-terminus hydrophobicity indicated by the Miyazawa score of the last 30 amino acids for high-confidence ORFs versus all predicted ORFs categorized by different start codons.

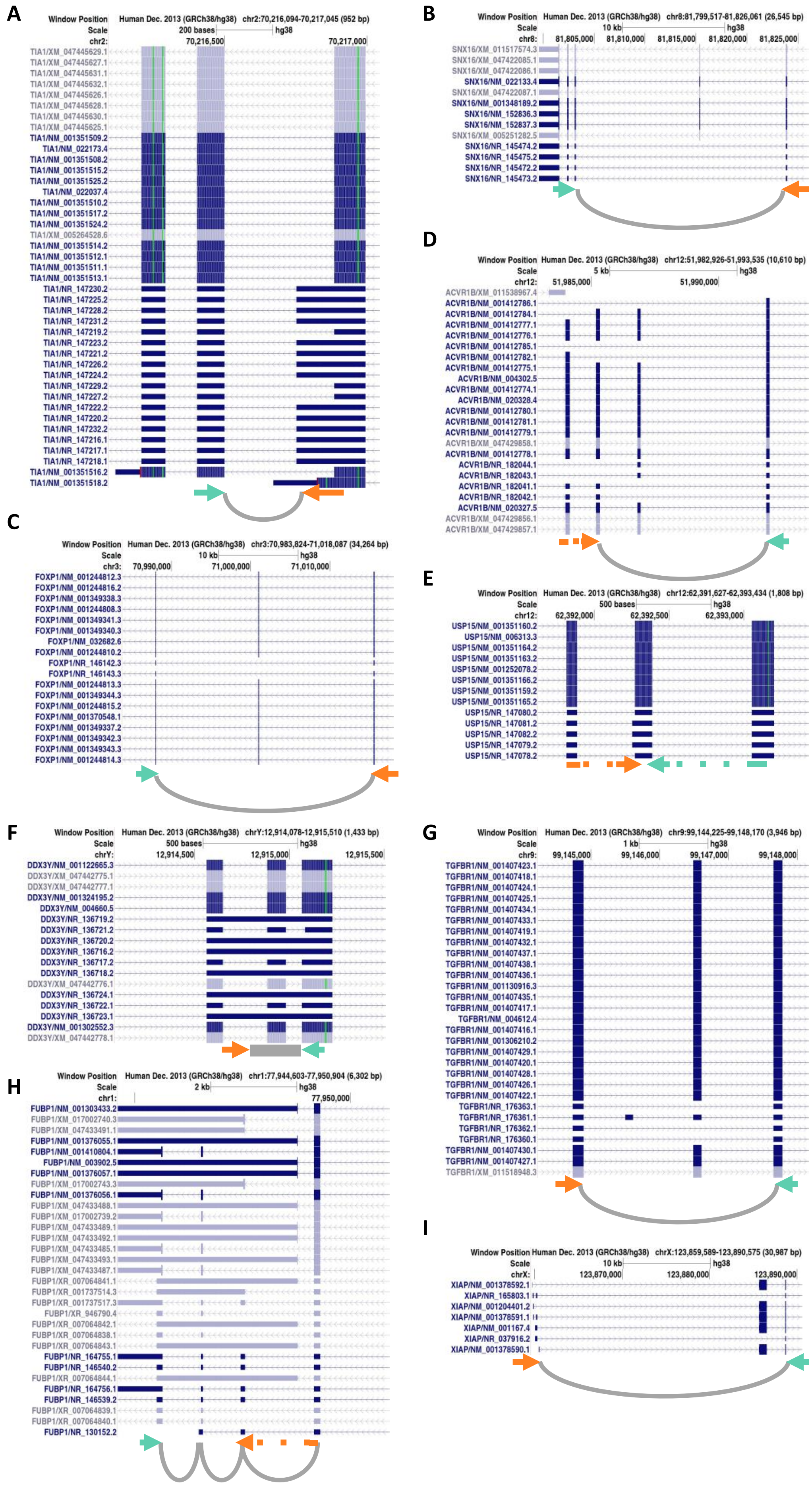


**Figure S10** shows the approach for primer design for 9 select bifunctional genes. The amplicon has been designed to either cover unique exon-exon junctions or extension of exons specific to non-coding transcripts; orange arrow represents the forward primer, and the green arrow represents the reverse primer. Dashes in either the forward or the reverse primer denote that the primer spans the specific exon-exon junction. The UCSC Genome Browser was used for this annotation representation.

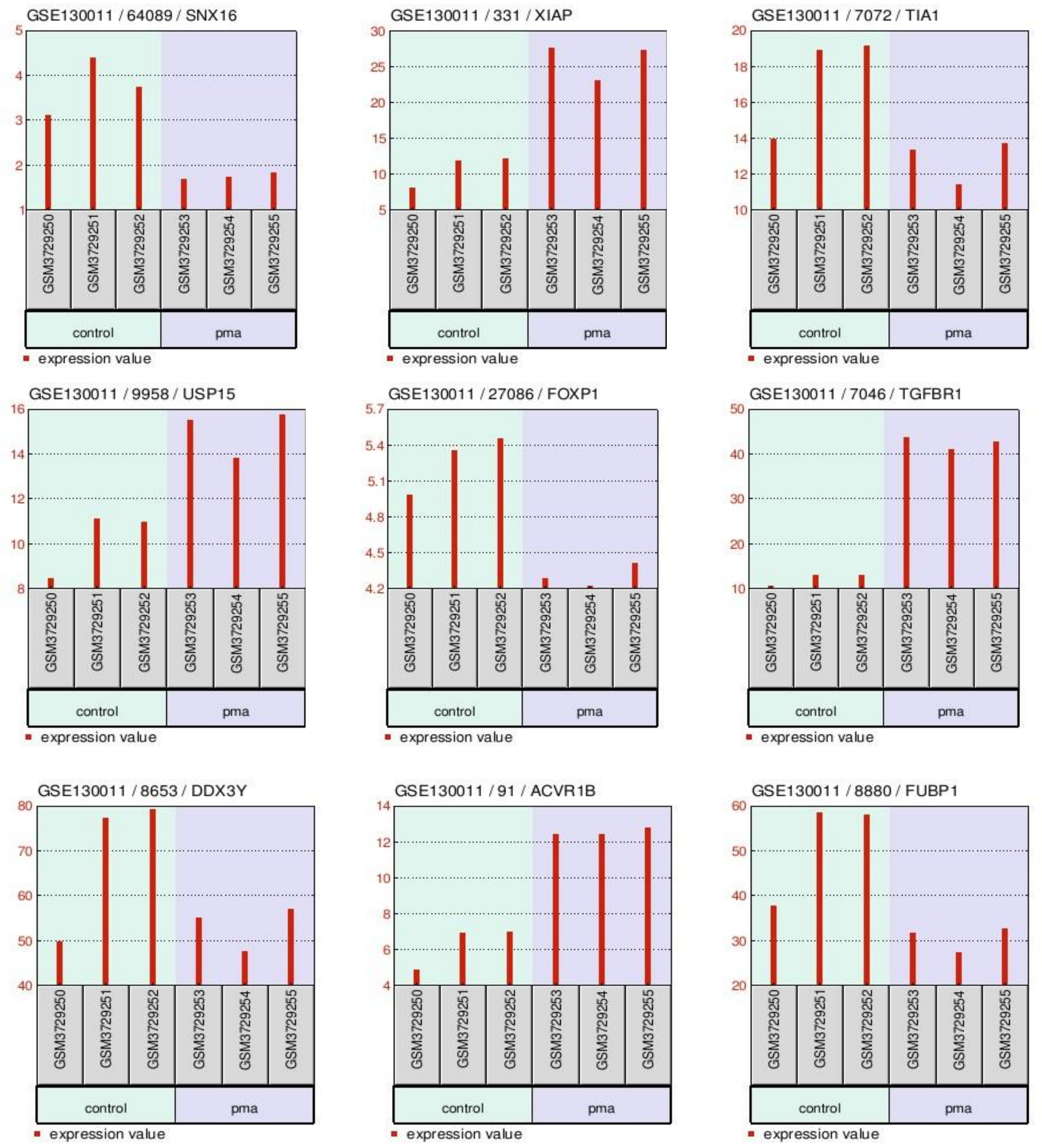


**Figure S11** shows the expression of selected bifunctional genes in THP1 cells with PMA differentiation from GSE130011 (https://www.ncbi.nlm.nih.gov/geo/query/acc.cgi?acc=GSE130011).

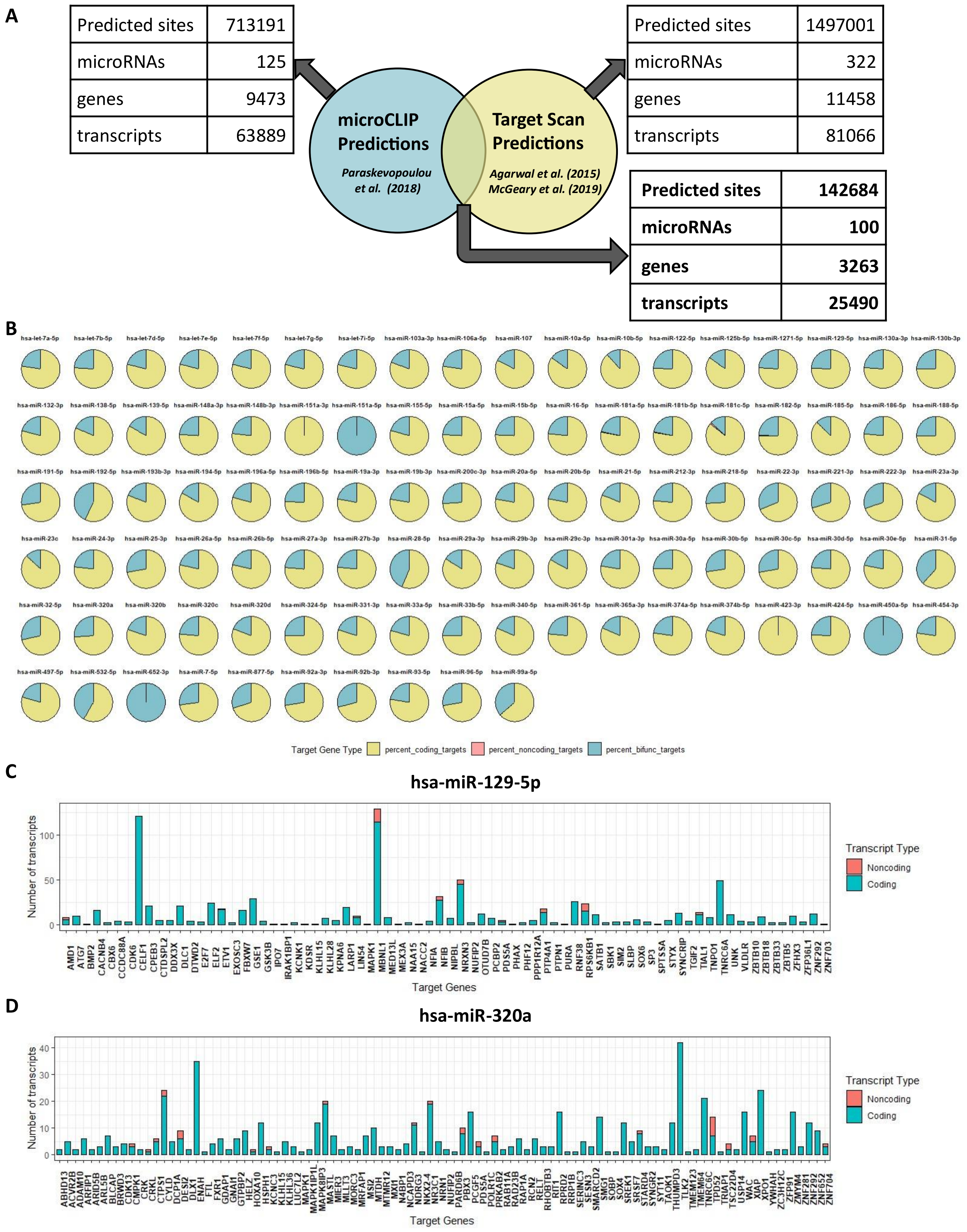


**Figure S12 (A)** Common miRNA binding sites from two predictions: microCLIP[3] and TargetScan[4], **(B)** miRNA binding targets include a significant number of bifunctional genes, as shown by the pie plots which represent the combined high-confidence miRNA-targets. An overview of target transcripts and their annotated biotypes for **(C)** hsa-miR-129-5p and **(D)** hsa-miR-320a.

**SUPPLEMENTAL TABLES**

**Table S1 Annotated and transcribed gene numbers from NCBI RefSeq GRCh38.p14 (GCF_000001405.40), related to Figure 1A** (number is number annotated and num_trans is number of annotated transcribed genes)

| gene_type | Number | number_trans | percent_num | percent_num_trans |
|---|---|---|---|---|
| transcribed_pseudogene | 1591 | 1241 | 0.023621 | 0.028869 |
| miRNA | 2139 | 1913 | 0.031757 | 0.044502 |
| lncRNA | 19645 | 18244 | 0.291659 | 0.424407 |
| Pseudogene | 17332 | 0 | 0.257319 | 0 |
| protein_coding | 23314 | 20048 | 0.346131 | 0.466374 |
| snRNA | 172 | 155 | 0.002554 | 0.003606 |
| snoRNA | 1300 | 1199 | 0.0193 | 0.027892 |
| ncRNA | 54 | 52 | 0.000802 | 0.00121 |
| antisense_RNA | 20 | 19 | 0.000297 | 0.000442 |
| tRNA | 691 | 0 | 0.010259 | 0 |
| ncRNA_pseudogene | 12 | 4 | 0.000178 | 9.31E-05 |
| V_segment | 365 | 0 | 0.005419 | 0 |
| misc_RNA | 68 | 44 | 0.00101 | 0.001024 |
| rRNA | 80 | 53 | 0.001188 | 0.001233 |
| Other | 29 | 0 | 0.000431 | 0 |
| C_region | 33 | 0 | 0.00049 | 0 |
| J_segment | 117 | 0 | 0.001737 | 0 |
| V_segment_pseudogene | 299 | 0 | 0.004439 | 0 |
| telomerase_RNA | 1 | 1 | 1.48E-05 | 2.33E-05 |
| vault_RNA | 4 | 4 | 5.94E-05 | 9.31E-05 |
| D_segment | 61 | 0 | 0.000906 | 0 |
| J_segment_pseudogene | 11 | 0 | 0.000163 | 0 |
| Y_RNA | 4 | 4 | 5.94E-05 | 9.31E-05 |
| RNase_MRP_RNA | 1 | 1 | 1.48E-05 | 2.33E-05 |
| scRNA | 4 | 4 | 5.94E-05 | 9.31E-05 |
| RNase_P_RNA | 2 | 1 | 2.97E-05 | 2.33E-05 |
| C_region_pseudogene | 7 | 0 | 0.000104 | 0 |

**Table S2 Pairwise comparisons of characteristics between Bifunctional genes and Coding/Noncoding genes, related to Figure 1D-F & Figure S1 C-E**

| Characteristic | Gene Type Comparison | RefSeq transcripts | p-value Welch's t-test | Cohen's d | Cohen's d interpretation |
|---|---|---|---|---|---|
| **Gene Length** | Bifunctional versus Coding | Complete | p < 2.2e-16 | 0.525 | moderate |
| | Bifunctional versus Noncoding | Complete | p < 2.2e-16 | 1.24 | large |
| **Number of Transcripts** | Bifunctional versus Coding | Complete | p < 2.2e-16 | 1.21 | large |
| | Bifunctional versus Noncoding | Complete | p < 2.2e-16 | 2.82 | large |
| **Number of exons** | Bifunctional versus Coding | Complete | p < 2.2e-16 | 0.619 | moderate |
| | Bifunctional versus Noncoding | Complete | p < 2.2e-16 | 2.31 | large |
| **Gene Length** | Bifunctional versus Coding | Validated transcripts | p < 2.2e-16 | 0.372 | small |
| | Bifunctional versus Noncoding | Validated transcripts | p < 2.2e-16 | 1.24 | large |
| **Number of Transcripts** | Bifunctional versus Coding | Validated transcripts | p < 2.2e-16 | 1.05 | large |
| | Bifunctional versus Noncoding | Validated transcripts | p < 2.2e-16 | 2.88 | large |
| **Number of exons** | Bifunctional versus Coding | Validated transcripts | p < 2.2e-16 | 0.454 | small |
| | Bifunctional versus Noncoding | Validated transcripts | p < 2.2e-16 | 2.08 | large |

**Table S3 Distribution of Coding, Noncoding and Bifunctional genes on chromosomes and strands, related to Figure 1G & 1H**

| chr | all_genes | Coding | Bifunctional | Noncoding | '+' strand (bifunctional genes) | '-' strand (bifunctional genes) |
|---|---|---|---|---|---|---|
| **chr1** | 4157 | 1647 | 418 | 2092 | 215 | 203 |
| **chr2** | 3009 | 915 | 339 | 1755 | 184 | 155 |
| **chr3** | 2371 | 796 | 270 | 1305 | 132 | 138 |
| **chr4** | 1909 | 610 | 161 | 1138 | 72 | 89 |
| **chr5** | 2096 | 717 | 187 | 1192 | 89 | 98 |
| **chr6** | 2327 | 816 | 236 | 1275 | 114 | 122 |
| **chr7** | 2104 | 724 | 213 | 1167 | 99 | 114 |
| **chr8** | 1770 | 548 | 166 | 1056 | 87 | 79 |
| **chr9** | 1799 | 596 | 183 | 1020 | 99 | 84 |
| **chr10** | 1772 | 536 | 198 | 1038 | 100 | 98 |
| **chr11** | 2335 | 1060 | 246 | 1029 | 126 | 120 |
| **chr12** | 2099 | 768 | 257 | 1074 | 122 | 135 |
| **chr13** | 1066 | 256 | 83 | 727 | 40 | 43 |
| **chr14** | 1385 | 484 | 124 | 777 | 64 | 60 |
| **chr15** | 1545 | 452 | 165 | 928 | 73 | 92 |
| **chr16** | 1682 | 666 | 191 | 825 | 98 | 93 |
| **chr17** | 2144 | 938 | 231 | 975 | 116 | 115 |
| **chr18** | 802 | 211 | 59 | 532 | 35 | 24 |
| **chr19** | 2204 | 1189 | 263 | 752 | 126 | 137 |
| **chr20** | 1140 | 432 | 105 | 603 | 59 | 46 |
| **chr21** | 636 | 196 | 43 | 397 | 19 | 24 |
| **chr22** | 928 | 353 | 99 | 476 | 53 | 46 |
| **chrX** | 1451 | 729 | 120 | 602 | 66 | 54 |
| **chrY** | 214 | 46 | 19 | 149 | 16 | 3 |
| **chrUn** | 42 | 4 | 2 | 36 | 1 | 1 |

**Table S4. Primer sequences used for real-time qPCR**

| Sl. No. | Species | Primer Name | Primer Sequence | Gene |
|---|---|---|---|---|
| 1 | Human | ACVR1B_NRqpcrFwd | CCTGGAATTGCTCATCGAGACT | ACVR1B |
| 2 | Human | ACVR1B_NRqpcrRev | GCTGATATTCTTCATGGACTCCGT | |
| 3 | Human | DDX3YNRqpcrFwd | ACTGCAGCATTTCTTTTACCCATAC | DDX3Y |
| 4 | Human | DDX3YNRqpcrRev | GTGTATCTGGCATCCGTTAGAAAAG | |
| 5 | Human | FOXP1_NRqpcrFwd | ACATGCCTCTACCAATGGACAG | FOXP1 |
| 6 | Human | FOXP1_NRqpcrRev | GCGTTCTTTGTCTTTTGCAAGTTT | |
| 7 | Human | FUBP1_NRqpcrFwd | GGACTGTGCTGTGTATCATCCT | FUBP1 |
| 8 | Human | FUBP1_NRqpcrRev | AGCTATGATTTGGTGCTTGCAG | |
| 9 | Human | SNX16NRqpcrFwd | GCTAACTGGCATTCTGTGAAACTT | SNX16 |
| 10 | Human | SNX16NRqpcrRev | CTGAGCAGTTTCTTTAGTGATTCCA | |
| 11 | Human | TGFBR1_NRqpcrFwd | GGCAGTAAGACATGATTCAGCC | TGFBR1 |
| 12 | Human | TGFBR1_NRqpcrRev | GCAGTTGGTAATCTTCATGAATTCCTT | |
| 13 | Human | TIA1NRqpcrFwd | TAGAGCTGTTCCCGGAGACTTA | TIA1 |
| 14 | Human | TIA1NRqpcrRev | CCCAGAAGTAACACCTCCACAG | |
| 15 | Human | USP15NRqpcrFwd | CTGCTGAGAAATACTTCTCTTTAGGAC | USP15 |
| 16 | Human | USP15NRqpcrRev | TACAATTCGGACAATACCAGGGATC | |
| 17 | Human | XIAP_NRqpcrFwd | CAGAGCGGAGTTGGCATTTC | XIAP |
| 18 | Human | XIAP_NRqpcrRev | AGCACTTTACTTTATCACCTTCACC | |
| 19 | Human | huCyclophilin_qPCR_F | GTCAACCCCACCGTGTTCTT | PPIA |
| 20 | Human | huCyclophilin_qPCR_R | CTGCTGTCTTTGGGACCTTGT | |

**Table S5 Expression of non-coding isoforms of bifunctional genes across cell lines**

| Cell Line | Gene | Rep1 | Rep2 | Rep3 | Average | stdev | p-value from two tailed Student t-test |
|---|---|---|---|---|---|---|---|
| **A549** | SNX16 | 1.00033 | 1.00264 | 1.00041 | 1.00113 | 0.00131 | |
| **PA1** | SNX16 | 5.50342 | 4.65721 | 5.73380 | 5.29814 | 0.56689 | 0.00019 |
| **PANC1** | SNX16 | 1.61901 | 1.20057 | 1.86680 | 1.56212 | 0.33674 | 0.04476 |
| **THP1** | SNX16 | 1.93164 | 1.56164 | 1.88877 | 1.79402 | 0.20238 | 0.00246 |
| **HT29** | SNX16 | 0.99421 | 1.05266 | 2.89567 | 1.64752 | 1.08133 | 0.35898 |
| **RD** | SNX16 | 3.20140 | 3.61171 | 4.18918 | 3.66743 | 0.49624 | 0.00074 |
| **A549** | XIAP | 1.15860 | 0.97909 | 1.68945 | 1.27571 | 0.36938 | |
| **PA1** | XIAP | 0.65638 | 1.66609 | 0.73174 | 1.01807 | 0.56247 | 0.54349 |
| **PANC1** | XIAP | 0.90860 | 0.84679 | 1.21668 | 0.99069 | 0.19814 | 0.30418 |
| **THP1** | XIAP | 0.31964 | 0.36907 | 0.44681 | 0.37851 | 0.06411 | 0.01432 |
| **HT29** | XIAP | 1.79575 | 1.38796 | 1.19290 | 1.45887 | 0.30762 | 0.54534 |
| **RD** | XIAP | 1.34296 | 0.97055 | 1.36513 | 1.22621 | 0.22168 | 0.85196 |
| **A549** | TIA1 | 1.00154 | 1.00070 | 1.00150 | 1.00124 | 0.00047 | |
| **PA1** | TIA1 | 0.79483 | 1.13991 | 1.69531 | 1.21001 | 0.45431 | 0.47064 |
| **PANC1** | TIA1 | 0.32153 | 0.52117 | 0.89670 | 0.57980 | 0.29203 | 0.06679 |
| **THP1** | TIA1 | 1.47817 | 1.27453 | 1.44996 | 1.40089 | 0.11033 | 0.00329 |
| **HT29** | TIA1 | 0.17831 | 0.15129 | 0.63607 | 0.32189 | 0.27243 | 0.01245 |
| **RD** | TIA1 | 0.83357 | 0.41908 | 0.36152 | 0.53805 | 0.25753 | 0.03569 |
| **A549** | USP15 | 1.01000 | 1.00409 | 1.03236 | 1.01548 | 0.01491 | |
| **PA1** | USP15 | 1.26466 | 1.80765 | 1.91543 | 1.66258 | 0.34879 | 0.03258 |
| **PANC1** | USP15 | 0.68295 | 0.72267 | 0.46232 | 0.62265 | 0.14026 | 0.00850 |
| **THP1** | USP15 | 3.04660 | 3.83780 | 1.57013 | 2.81817 | 1.15096 | 0.05339 |
| **HT29** | USP15 | 0.94448 | 1.05052 | 1.58934 | 1.19478 | 0.34579 | 0.42030 |
| **RD** | USP15 | 1.73104 | 1.73701 | 0.99375 | 1.48727 | 0.42741 | 0.12864 |
| **A549** | FOXP1 | 1.01812 | 1.01561 | 1.26685 | 1.10019 | 0.14433 | |
| **PA1** | FOXP1 | 3.60984 | 1.13169 | 0.66256 | 1.80136 | 1.58365 | 0.48760 |
| **PANC1** | FOXP1 | 1.04444 | 0.52949 | 0.70358 | 0.75917 | 0.26194 | 0.11948 |
| **HT29** | FOXP1 | 4.63916 | 1.82245 | 4.85716 | 3.77292 | 1.69268 | 0.05271 |
| **RD** | FOXP1 | 5.53738 | 2.57240 | 0.98086 | 3.03021 | 2.31250 | 0.22256 |
| **THP1** | FOXP1 | 4.52302 | 4.86355 | 4.51581 | 4.63413 | 0.19872 | 0.00002 |
| **A549** | TGFBR1 | 1.01924 | 1.08472 | 1.00595 | 1.03664 | 0.04217 | |
| **PA1** | TGFBR1 | 0.59594 | 0.41389 | 0.61222 | 0.54068 | 0.11011 | 0.00189 |
| **PANC1** | TGFBR1 | 0.30439 | 0.29359 | 0.43725 | 0.34508 | 0.08001 | 0.00019 |
| **THP1** | TGFBR1 | 1.46121 | 0.61915 | 1.23361 | 1.10466 | 0.43559 | 0.80109 |
| **HT29** | TGFBR1 | 0.30245 | 0.45843 | 0.74359 | 0.50149 | 0.22370 | 0.01520 |
| **RD** | TGFBR1 | 1.71571 | 1.06538 | 2.37848 | 1.71986 | 0.65656 | 0.14646 |
| **A549** | DDX3Y | 1.00709 | 0.98703 | 1.00417 | 0.99943 | 0.01084 | |
| **PA1** | DDX3Y | 0.00339 | 0.00000 | 0.00000 | 0.00113 | 0.00196 | 0.00000 |
| **PANC1** | DDX3Y | 0.00000 | 0.00000 | 0.00000 | 0.00000 | 0.00000 | 0.00000 |
| **THP1** | DDX3Y | 0.46523 | 0.62417 | 0.44928 | 0.51289 | 0.09670 | 0.00098 |
| **HT29** | DDX3Y | 0.00000 | 0.00000 | 0.00000 | 0.00000 | 0.00000 | 0.00000 |
| **RD** | DDX3Y | 0.00000 | 0.00000 | 0.00155 | 0.00052 | 0.00090 | 0.00000 |
| **A549** | ACVR1B | 1.00600 | 0.99994 | 1.00975 | 1.00523 | 0.00495 | |
| **PA1** | ACVR1B | 1.45477 | 2.74993 | 1.75570 | 1.98680 | 0.67780 | 0.06618 |
| **PANC1** | ACVR1B | 0.47340 | 0.50278 | 0.79172 | 0.58930 | 0.17592 | 0.01493 |
| **THP1** | ACVR1B | 0.42831 | 1.11545 | 1.00954 | 0.85110 | 0.36996 | 0.51051 |
| **HT29** | ACVR1B | 0.48973 | 0.76127 | 1.05911 | 0.77004 | 0.28479 | 0.22590 |
| **RD** | ACVR1B | 0.73562 | 1.27493 | 0.80310 | 0.93788 | 0.29383 | 0.71169 |

| | | | | | | | |
|---|---|---|---|---|---|---|---|
| **A549** | FUBP1 | 1.01351 | 0.99149 | 1.00024 | 1.00174 | 0.01109 | |
| **PA1** | FUBP1 | 12.59931 | 7.67696 | 6.52867 | 8.93498 | 3.22492 | 0.01305 |
| **PANC1** | FUBP1 | 0.91815 | 0.52341 | 0.66239 | 0.70132 | 0.20023 | 0.06037 |
| **THP1** | FUBP1 | 7.55288 | 3.02254 | 3.69127 | 4.75556 | 2.44551 | 0.05647 |
| **HT29** | FUBP1 | 0.87532 | 0.81253 | 1.97577 | 1.22120 | 0.65423 | 0.59245 |
| **RD** | FUBP1 | 4.74549 | 2.74346 | 3.05013 | 3.51303 | 1.07830 | 0.01569 |

**Table S6 Expression of non-coding isoforms during THP1 differentiation and inflammation induction with LPS**

| Condition | Gene | Rep1 | Rep2 | Rep3 | Average | stdev | p-value from two tailed Student t-test |
|---|---|---|---|---|---|---|---|
| **THP1** | SNX16 | 1.00424 | 1.00004 | 1.00068 | 1.00165 | 0.00226 | |
| **THP1+PMA** | SNX16 | 0.83482 | 1.01924 | 0.40392 | 0.75266 | 0.31578 | 0.24379 |
| **THP1+PMA+LPS** | SNX16 | 1.26316 | 1.36020 | 0.72734 | 1.11690 | 0.34084 | 0.58957 |
| **THP1** | XIAP | 1.00069 | 1.00819 | 1.00363 | 1.00417 | 0.00378 | |
| **THP1+PMA** | XIAP | 8.71360 | 5.11011 | 2.89878 | 5.57416 | 2.93505 | 0.05427 |
| **THP1+PMA+LPS** | XIAP | 4.94661 | 3.58844 | 3.74043 | 4.09183 | 0.74415 | 0.00199 |
| **THP1** | TIA1 | 1.00099 | 1.00680 | 1.00680 | 1.00486 | 0.00335 | |
| **THP1+PMA** | TIA1 | 0.60955 | 0.35417 | 0.20897 | 0.39090 | 0.20280 | 0.00633 |
| **THP1+PMA+LPS** | TIA1 | 0.43745 | 0.25723 | 0.43169 | 0.37546 | 0.10243 | 0.00044 |
| **THP1** | USP15 | 1.00129 | 1.00903 | 1.00026 | 1.00353 | 0.00480 | |
| **THP1+PMA** | USP15 | 2.35404 | 2.17429 | 1.24120 | 1.92318 | 0.59741 | 0.05602 |
| **THP1+PMA+LPS** | USP15 | 2.46229 | 2.38508 | 2.11265 | 2.32001 | 0.18368 | 0.00024 |
| **THP1** | FOXP1 | 1.00743 | 1.00406 | 1.02030 | 1.01059 | 0.00857 | |
| **THP1+PMA** | FOXP1 | 0.54598 | 0.25041 | 0.17468 | 0.32369 | 0.19620 | 0.00375 |
| **THP1+PMA+LPS** | FOXP1 | 0.69134 | 0.55961 | 0.77708 | 0.67601 | 0.10954 | 0.00619 |
| **THP1** | FUBP1 | 1.00174 | 1.00233 | 1.01586 | 1.00665 | 0.00799 | |
| **THP1+PMA** | FUBP1 | 0.06234 | 0.04662 | 0.03724 | 0.04873 | 0.01268 | 0.00000 |
| **THP1+PMA+LPS** | FUBP1 | 0.08015 | 0.10581 | 0.19501 | 0.12699 | 0.06029 | 0.00002 |
| **THP1** | TGFBR1 | 1.00434 | 1.01003 | 1.00143 | 1.00527 | 0.00438 | |
| **THP1+PMA** | TGFBR1 | 1.16896 | 1.89829 | 0.42745 | 1.16490 | 0.73543 | 0.72604 |
| **THP1+PMA+LPS** | TGFBR1 | 1.46269 | 1.20504 | 0.79833 | 1.15535 | 0.33495 | 0.48105 |
| **THP1** | ACVR1B | 1.01118 | 1.00972 | 1.05087 | 1.02392 | 0.02334 | |
| **THP1+PMA** | ACVR1B | 1.77278 | 1.70830 | 0.48893 | 1.32334 | 0.72334 | 0.51324 |
| **THP1+PMA+LPS** | ACVR1B | 1.01215 | 0.43447 | 0.61845 | 0.68836 | 0.29512 | 0.12109 |
| **THP1** | DDX3Y | 1.01897 | 1.00667 | 1.02620 | 1.01728 | 0.00987 | |
| **THP1+PMA** | DDX3Y | 0.67526 | 0.52382 | 0.28606 | 0.49505 | 0.19619 | 0.01000 |
| **THP1+PMA+LPS** | DDX3Y | 1.08867 | 0.90798 | 2.36201 | 1.45289 | 0.79249 | 0.39502 |

**Table S7 TPM Sums for RefSeq annotated transcrtipts from isoQuant on direct cDNA sequencing samples from SGNEx, related to Figure S6**

| S. No. | Gene Type | Sample | Sum of TPM | Cell Line |
|---|---|---|---|---|
| 1 | Bifunctional | A549_rep1_run3 | 140905.8369 | A549 |
| 2 | Coding | A549_rep1_run3 | 818537.0497 | A549 |
| 3 | Noncoding | A549_rep1_run3 | 23994.1809 | A549 |
| 4 | Bifunctional | A549_rep2_run1 | 122890.6887 | A549 |
| 5 | Coding | A549_rep2_run1 | 807946.4456 | A549 |
| 6 | Noncoding | A549_rep2_run1 | 39836.28449 | A549 |
| 7 | Bifunctional | A549_rep3_run2 | 135433.3054 | A549 |
| 8 | Coding | A549_rep3_run2 | 787314.6893 | A549 |
| 9 | Noncoding | A549_rep3_run2 | 31868.69167 | A549 |
| 10 | Bifunctional | A549_rep5_run3 | 123677.857 | A549 |
| 11 | Coding | A549_rep5_run3 | 818349.0458 | A549 |
| 12 | Noncoding | A549_rep5_run3 | 30345.41785 | A549 |
| 13 | Bifunctional | H9_rep2_run3 | 132719.7537 | H9 |
| 14 | Coding | H9_rep2_run3 | 764727.4649 | H9 |
| 15 | Noncoding | H9_rep2_run3 | 46447.56071 | H9 |
| 16 | Bifunctional | H9_rep3_run3 | 120431.146 | H9 |
| 17 | Coding | H9_rep3_run3 | 759574.7545 | H9 |
| 18 | Noncoding | H9_rep3_run3 | 50128.32187 | H9 |
| 19 | Bifunctional | H9_rep4_run3 | 119418.0711 | H9 |
| 20 | Coding | H9_rep4_run3 | 753612.9665 | H9 |
| 21 | Noncoding | H9_rep4_run3 | 53188.60922 | H9 |
| 22 | Bifunctional | HEYA8_rep1_run3 | 122773.4728 | HEYA8 |
| 23 | Coding | HEYA8_rep1_run3 | 759323.1329 | HEYA8 |
| 24 | Noncoding | HEYA8_rep1_run3 | 49666.52466 | HEYA8 |
| 25 | Bifunctional | HEYA8_rep2_run3 | 112779.7742 | HEYA8 |
| 26 | Coding | HEYA8_rep2_run3 | 815534.2459 | HEYA8 |
| 27 | Noncoding | HEYA8_rep2_run3 | 38001.82528 | HEYA8 |
| 28 | Bifunctional | HEYA8_rep3_run2 | 113591.9671 | HEYA8 |
| 29 | Coding | HEYA8_rep3_run2 | 813118.3943 | HEYA8 |
| 30 | Noncoding | HEYA8_rep3_run2 | 39273.0303 | HEYA8 |
| 31 | Bifunctional | HCT116_rep1_run4 | 113954.8641 | HCT116 |
| 32 | Coding | HCT116_rep1_run4 | 817617.2638 | HCT116 |
| 33 | Noncoding | HCT116_rep1_run4 | 36222.67342 | HCT116 |
| 34 | Bifunctional | HCT116_rep3_run2 | 31484.00432 | HCT116 |
| 35 | Coding | HCT116_rep3_run2 | 188518.8143 | HCT116 |
| 36 | Noncoding | HCT116_rep3_run2 | 770893.0333 | HCT116 |
| 37 | Bifunctional | HCT116_rep4_run1 | 141775.7974 | HCT116 |
| 38 | Coding | HCT116_rep4_run1 | 794299.6311 | HCT116 |
| 39 | Noncoding | HCT116_rep4_run1 | 27246.42633 | HCT116 |
| 40 | Bifunctional | HCT116_rep5_run1 | 138726.44 | HCT116 |
| 41 | Coding | HCT116_rep5_run1 | 794636.0277 | HCT116 |
| 42 | Noncoding | HCT116_rep5_run1 | 27976.51885 | HCT116 |
| 43 | Bifunctional | HEPG2_rep1_run1 | 140621.4804 | HEPG2 |
| 44 | Coding | HEPG2_rep1_run1 | 789245.9238 | HEPG2 |
| 45 | Noncoding | HEPG2_rep1_run1 | 28351.49793 | HEPG2 |
| 46 | Bifunctional | HEPG2_rep1_run2 | 102600.5112 | HEPG2 |
| 47 | Coding | HEPG2_rep1_run2 | 778875.4108 | HEPG2 |
| 48 | Noncoding | HEPG2_rep1_run2 | 62294.61641 | HEPG2 |

| | | | | |
|---|---|---|---|---|
| **49** | Bifunctional | HEPG2_rep2_run1 | 109749.4292 | HEPG2 |
| **50** | Coding | HEPG2_rep2_run1 | 786010.9445 | HEPG2 |
| **51** | Noncoding | HEPG2_rep2_run1 | 56049.7218 | HEPG2 |
| **52** | Bifunctional | HEPG2_rep4_run1 | 108360.17 | HEPG2 |
| **53** | Coding | HEPG2_rep4_run1 | 808476.1046 | HEPG2 |
| **54** | Noncoding | HEPG2_rep4_run1 | 44183.9495 | HEPG2 |
| **55** | Bifunctional | HEPG2_rep4_run2 | 105798.5292 | HEPG2 |
| **56** | Coding | HEPG2_rep4_run2 | 793359.1453 | HEPG2 |
| **57** | Noncoding | HEPG2_rep4_run2 | 58374.76231 | HEPG2 |
| **58** | Bifunctional | HEPG2_rep5_run3 | 113978.883 | HEPG2 |
| **59** | Coding | HEPG2_rep5_run3 | 807339.8229 | HEPG2 |
| **60** | Noncoding | HEPG2_rep5_run3 | 37909.86995 | HEPG2 |
| **61** | Bifunctional | K562_rep1_run2 | 104705.0943 | K562 |
| **62** | Coding | K562_rep1_run2 | 757388.2132 | K562 |
| **63** | Noncoding | K562_rep1_run2 | 58083.8399 | K562 |
| **64** | Bifunctional | K562_rep2_run1 | 93743.657 | K562 |
| **65** | Coding | K562_rep2_run1 | 760129.2801 | K562 |
| **66** | Noncoding | K562_rep2_run1 | 78067.66906 | K562 |
| **67** | Bifunctional | K562_rep3_run1 | 91228.58011 | K562 |
| **68** | Coding | K562_rep3_run1 | 794639.908 | K562 |
| **69** | Noncoding | K562_rep3_run1 | 57148.33067 | K562 |
| **70** | Bifunctional | K562_rep4_run2 | 102503.2974 | K562 |
| **71** | Coding | K562_rep4_run2 | 794005.6974 | K562 |
| **72** | Noncoding | K562_rep4_run2 | 41912.8408 | K562 |
| **73** | Bifunctional | MCF7_rep1_run2 | 136823.6107 | MCF7 |
| **74** | Coding | MCF7_rep1_run2 | 748276.6235 | MCF7 |
| **75** | Noncoding | MCF7_rep1_run2 | 26791.54345 | MCF7 |
| **76** | Bifunctional | MCF7_rep3_run3 | 90841.94978 | MCF7 |
| **77** | Coding | MCF7_rep3_run3 | 758692.3412 | MCF7 |
| **78** | Noncoding | MCF7_rep3_run3 | 48857.34841 | MCF7 |
| **79** | Bifunctional | MCF7_rep4_run2 | 117989.5964 | MCF7 |
| **80** | Coding | MCF7_rep4_run2 | 753463.2144 | MCF7 |
| **81** | Noncoding | MCF7_rep4_run2 | 40380.49111 | MCF7 |
